# Human mitochondrial RNA polymerase structures reveal transcription start-site and slippage mechanism

**DOI:** 10.1101/2024.12.02.626445

**Authors:** Jiayu Shen, Quinten Goovaerts, Yogeeshwar Ajjugal, Brent De Wijngaert, Kalyan Das, Smita S. Patel

## Abstract

Human mitochondrial RNA polymerase (POLRMT) and protein factors TFAM and TFB2M assemble on mitochondrial DNA promoters to initiate promoter-specific transcription. We present cryo-EM structures of two initiation complexes, IC3 and slipped-IC3, with fully resolved transcription bubbles containing RNA transcripts starting from +1 and −1 positions, respectively. These structures reveal the mechanisms of promoter melting, start site selection, and slippage synthesis. Promoter melting begins at −4 with base-specific interactions of −4 and −3 template guanines with POLRMT and −1 non-template adenine with TFB2M, stabilizing the bubble and facilitating initiation from +1. Slippage occurs when a synthesized 2-mer RNA shifts to −1; the −1 position is not an alternative start-site. The conserved non-template sequence (-1)AAA(+2) is recognized by a non-template stabilizing loop (K_153_LDPRSGGVIKPP_165_) and Y209 from TFB2M and W1026 of POLRMT. The initiation complex on cryo-EM grids exist in equilibrium with apo and dimeric POLRMTs, whose relative concentrations may regulate transcription initiation.

**HIGHLIGHTS:**

- Cryo-EM structures of active human mitochondrial transcription initiation complexes
- De novo RNA synthesis begins at +1, linking +1 and +2 NTPs
- Synthesized 2-mer RNA anneals at −1, initiating RNA slippage synthesis
- POLRMT fingers’ motion alters TFB2M interactions with non-template strand

## INTRODUCTION

Transcription initiation is a critical step in the regulation of gene expression. During this multistep process, an RNA polymerase (RNAP) is recruited to a specific promoter site via interactions with DNA sequences and transcription factors. The RNAP initiation complex melts the promoter DNA between defined positions and initiates RNA synthesis at a transcription start site (TSS). In human mitochondria, transcription of mitochondrial DNA is carried out by a nuclear-encoded RNA polymerase, POLRMT, in coordination with two transcription factors, mitochondrial transcription factor A (TFAM) and mitochondrial transcription factor B2 (TFB2M)^1–4^. Human mitochondrial (h-mt) DNA contains three promoters, located on the light (L) strand (LSP and LSP2) and heavy (H) strand (HSP)^5–7^. The polycistronic RNAs from these promoters code for essential OXPHOS mRNA transcripts, mitochondrial ribosomal RNAs, and tRNAs^1^. Transcription from LSP also generates primers for initiating h-mt DNA replication^8,9^. We lack structures of the initiation complex intermediates of POLRMT. Understanding the physical basis of transcription initiation is crucial for elucidating the regulation of mitochondrial DNA maintenance and gene expression. The structural basis of initiation provides the foundation to address mitochondrial disorders from inherited mutations in POLRMT^1,10^, aid toxicity studies of antivirals^11^, and develop POLRMT- targeted anticancer therapies^12^.

POLRMT shares homology with the well-characterized single-subunit RNAPs from bacteriophage T7 and yeast mitochondria (y-mt) Rpo41^13,14^. Unlike T7 RNAP, which independently carries out all steps of transcription initiation^15,16^, y-mt RNAP and h-mt POLRMT require the transcription factor Mtf1 and TFB2M, respectively, for promoter melting^17–20^. T7 and y-mt promoters have extended regions of conserved DNA sequences^21,22^ that involve in promoter-specific interactions with protein residues to assist DNA melting^18,23,24^. In yeast, Mtf1 interacts with the non-template −4 to −2 nucleotides upstream of the start site in a sequence-specific manner, and these interactions remain intact during RNA synthesis from 2 to 8 nucleotides^25,26^. In humans, TFB2M also interacts with the non-template strand^27^, however, the sequence conservation in the h-mt promoters is near the start site between −1 and +4^7^. Therefore, how TFB2M recognizes the non-template strand to orchestrate promoter melting remains unclear. Additionally, the initiation bubble size or the transcription start sites on h-mt promoters are not clearly defined. Mutations in the h-mt promoter between -5 and +3 affect transcription activity^7,28^, suggesting that POLRMT and TFB2M engage with the initiation bubble in a sequence-specific manner. However, the crystal structure of the open complex failed to resolve most of the initiation bubble and was captured in an inactive finger-clenched conformation, without RNA or NTPs bound^27^. Consequently, the promoter-melting mechanism and the precise structural elements of POLRMT and TFB2M involved in promoter melting and bubble stabilization during transcription initiation remain elusive.

Transcription initiation from LSP or HSP does not occur from a specific position, unlike the initiation observed on y-mt and T7 promoters. RNA-sequencing has mapped the 5’ ends of the transcripts from LSP and HSP to positions −1 and +1, with fewer reads mapping to position +2^5,7,28,29^. A recent report proposed that the RNA product starting from −1 TSS, which was historically assigned as +1, resulted from RNA slippage^28^. Both LSP and HSP have runs of T’s at their TSS that appear to be responsible for slippage; in bacterial and T7 RNAPs, slippage synthesis is known to occur when there are consecutive runs of T’s or G’s in the template sequences^30,31^. Repeated slippage synthesis results in transcripts containing a characteristic sequence of non-complementary A’s at the 5’-end of RNA, as observed in HSP transcripts^5,19,28,32^. However, this repeated slippage synthesis is not observed on LSP, where two distinct RNA transcripts are produced that differ by only one nucleotide in length. Structural basis of how these two RNA products are synthesized at the LSP will elucidate the mechanism for TSS selection and RNA slippage synthesis, including if both +1 and −1 positions can function as TSS.

In this study, we combined mutagenesis, biochemical assays, and cryo-EM structural studies to investigate the mechanism of transcription initiation at the LSP promoter. We report the structures of two h-mt RNAP initiation complexes (ICs)—IC3 and slipped IC3—capturing the synthesis of two distinct RNA transcripts. The transcripts, pppGp(3’-d)A and pppGpGpA, had their 5’-ends positioned at +1 and −1 in IC3 and slipped IC3 structures, respectively. This study reveals i) fully resolved transcription initiation bubbles that define the exact size of the initially melted regions in LSP, ii) two different RNA:DNA hybrid intermediates that help establish the transcription start site and provide the basis for RNA slippage, and iii) novel elements of TFB2M (NT-stabilizing loop and an adjacent helix) involved in non-template strand recognition, promoter melting, and start site selection. Our comprehensive structural and mechanistic studies provide a detailed depiction of the molecular machine initiating h-mt DNA transcription and offer a comparison with the analogous machinery in yeast mitochondria. Distinct interactions between the non-template strand and transcription factors TFB2M vs. Mtf1 in h-mt vs. y-mt ICs, respectively, differentiate their precise mechanisms of promoter melting and transcription initiation.

## RESULTS

### Nucleotide specificity requirement for transcription initiation

To design a minimal transcription bubble with an optimized sequence for cryo-EM studies, we conducted a systematic base scanning study of the transcription initiation region in LSP. Since mutation of the -5 base-pair (newly annotated numbering) is deleterious to transcription initiation^7,19,28^, we chose to create an LSP scaffold with a bubble from −4 to +1. The individual nucleotides in this region were mutated and their effects were assessed using runoff transcription assays on a 63-bp (-43 to +20) LSP fragment (Figure 1A) with purified POLRMT, TFB2M, and TFAM proteins. The wt LSP produces two runoff products: 18-nt and 19-nt in the presence of ATP, GTP, UTP, and 3’-dCTP (Figure 1B); the terminator 3’-dCTP prevents random nucleotide addition at the 3’-end of transcript. Thus, the 18-nt RNA product is synthesized from +1 TSS and the 19-nt product results from the −1 TSS or one-nucleotide slipped RNA.

**Figure 1.**
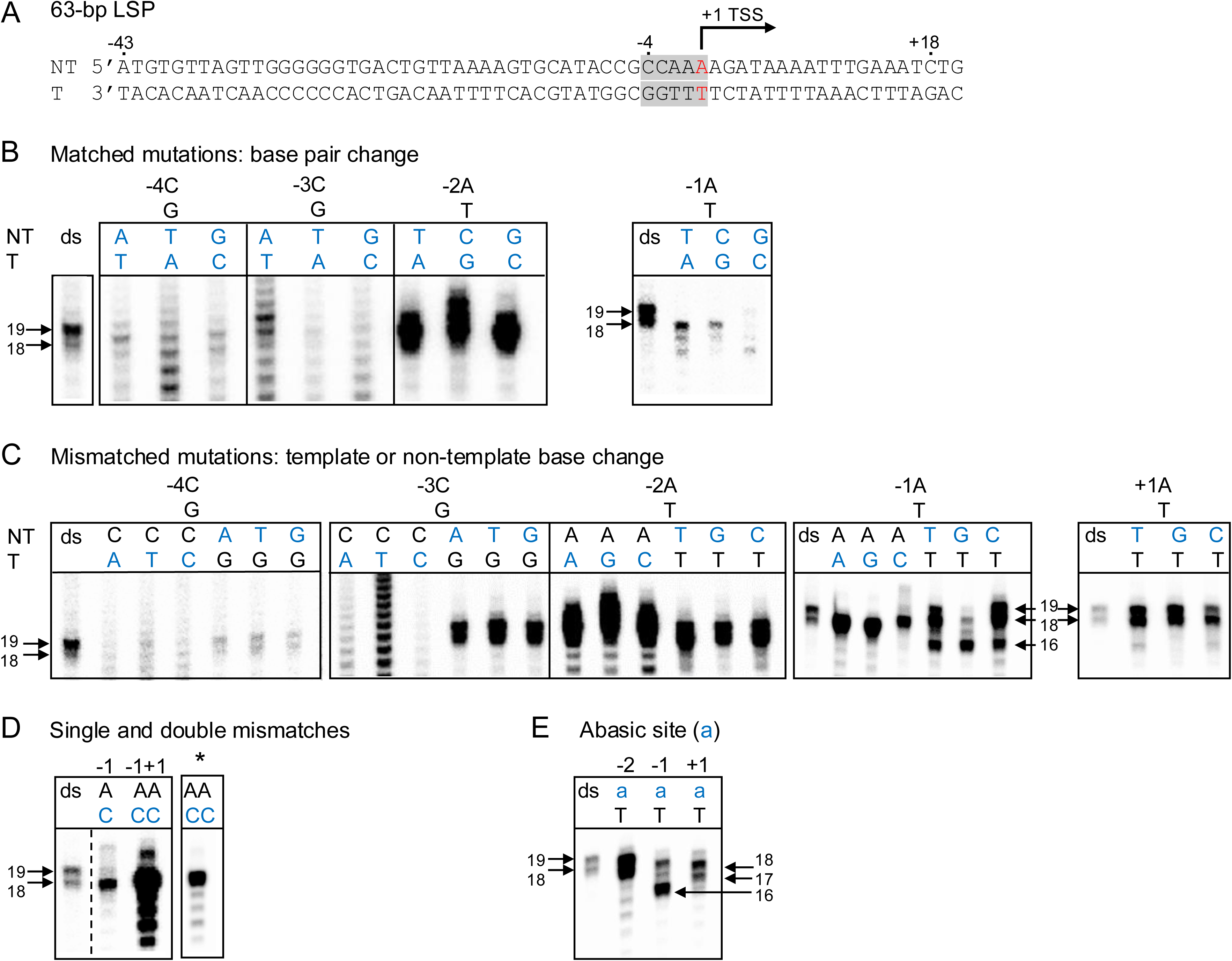
Nucleotide specificity requirement for transcription initiation. (A) The LSP DNA fragment used for the promoter mutational studies. The +1 TSS (newly annotated nomenclature) is in red, and the mutation positions, −4 to +1, are shaded gray. (B-E) Transcription runoff products from LSP with base-pair or base mutations in the −4 to +1 region. The mutated bases are shown in blue. In panel D, the lane with the asterisk is the same as the preceding lane but at a lower contrast to show the runoff product length. “ds” represents the duplex LSP without mismatches. Representative images from duplicates are shown.

We started by mutating each base pair from −4 to −1 to every possible alternative (Figure 1B). Any alteration of the −4 C:G or −3 C:G to other base pairs inhibited the specific runoff products and resulted in a ladder of RNAs presumably from non-specific transcription initiations. In contrast, changing the −2 A:T to other base pairs was well tolerated. Interestingly, mutating the - 1 A:T base pair produced shorter 17-nt and 16-nt RNA products, eliminated the 19-nt product, and significantly reduced the yield of the 18-nt product. These results indicate that the −4 to −1 promoter region has base-specific interactions with POLRMT and TFB2M as previously indicated^7,28^.

To pinpoint which specific template or non-template base is involved in base-specific interactions, we carried out a systematic promoter mutational study where each base between −4 to +1 was altered, resulting in six possible mismatched base pair combinations at each position that were tested in the transcription assay (Figure 1C). In general, when the mutated base did not disrupt specific interactions with the protein, we observed higher runoff product levels than the canonical base-paired LSP. This could be because the mismatched base pair facilitates promoter melting.

#### -4 and −3 template bases are crucial

Mutation of the template strand −4G or −3G to A, T, or C severely inhibited runoff synthesis (Figure 1C). In contrast, mutation of the non-template strand −4C or −3C had a lesser impact. The non-template −4C mutation decreased runoff synthesis but mutating the −3C led to increased synthesis likely from aiding promoter melting. Thus, template −4G and −3G are critical for transcription initiation.

#### -2 base changes are tolerated

Mutation of the −2 template T or non-template A had no deleterious effect on transcription and the mismatched promoters showed hyperactive runoff synthesis (Figure 1C).

#### -1 template base mutation inhibits the 19-nt product

Mutating the template −1T to A, C, or G eliminated the 19-nt product, resulting in only the 18-nt product (Figure 1C). This could be due to inefficient initiation from GTP or pyrimidines or a lack of RNA slippage due to interruption of the string of three T’s. To distinguish between the two events, we changed −1 and +1 template T’s to C’s while keeping the non-template A’s intact. The A:C mismatches provided equal opportunities for initiation with GTP from the −1 and +1 sites; however, only the +1 TSS 18-nt RNA was produced (Figure 1D). These results indicate that transcription can initiate from GTP, but only +1 TSS is active. Hence, the 19-nt product is from slippage synthesis^28^.

#### -1 non-template base is critical for defining the start site

Mutation of the non-template −1A to T, C, or G altered the transcription initiation profile (Figure 1C). A non-template −1T or C produced the 18- and 19-nt RNAs, but there was a predominant synthesis of a shorter 16-nt product, likely arising from the +3 TSS. This 16-nt RNA was overproduced when the non-template −1A was changed to G. Thus, the non-template −1 A has a critical role in transcription start site selection.

#### +1 non-template base variation is tolerated

Unlike the non-template −1A, mutation of +1A had no deleterious effect on the normal runoff RNA products (Figure 1C). The +1 non-template mismatches produced both 18- and 19-nt RNA runoffs at higher yields than the duplex LSP sequence.

The −1 and +1 non-template bases are important for slippage synthesis: To further test the roles of non-template bases, we substituted individual positions between −2 and +1 with abasic nucleotide (Figure 1E). The promoter with −2 abasic site had no deleterious effect and overproduced the 18-nt and 19-nt runoff RNAs. In comparison, substituting the −1A with abasic nucleotide drastically inhibited the 18-nt and 19-nt RNA products, and most of the initiation was from the +3 position, resulting in a predominant 16-nt product. This is consistent with the −1A’s role in TSS selection (Figure 1C). Mutation of the +1A to abasic nucleotide generated the RNA from +1 TSS (18-nt), and an additional product from +2 TSS (17-nt) but inhibited RNA slippage (19-nt). Thus, interactions of +1 nucleotide base support RNA slippage.

### Cryo-EM structures of the h-mt initiation complex IC3

An LSP bubble with mismatches in the −4 to +1 region was designed for cryo-EM studies by keeping the essential template and non-template bases intact, including template −4G and −3G, and non-template −1A and +1A (Figure 2A, Figure S1A). The RNA initiation region contained −1C and +1C template nucleotides which provide an opportunity to capture the IC intermediates from the two modes of RNA synthesis, from −1 and +1 positions, using appropriate initiating nucleotides. *In vitro* transcription assays confirmed the bubble promoter is active (Figures S1A-1C).

**Figure 2.**
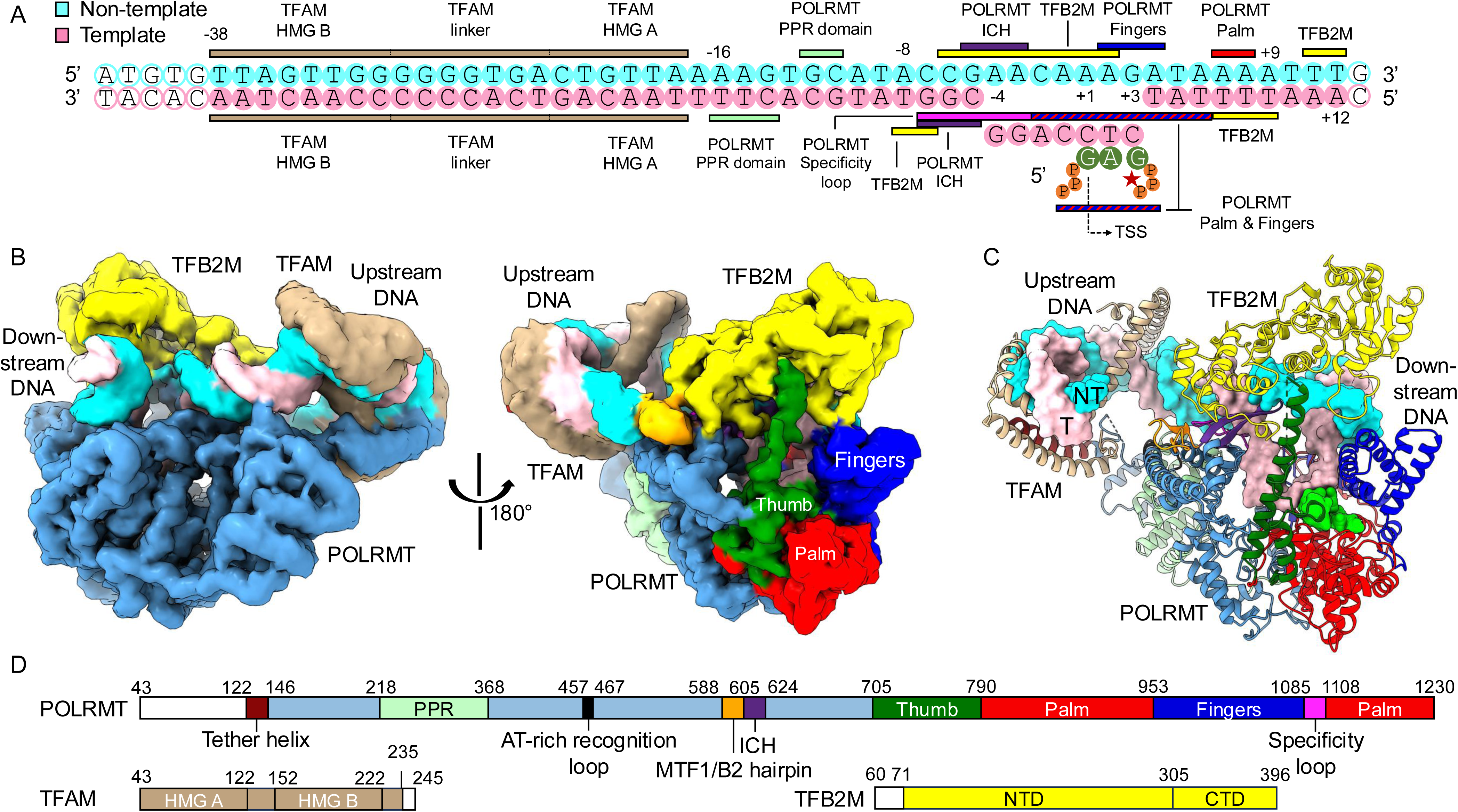
Structure of the human mitochondrial RNA polymerase transcription initiation complex with 2-mer RNA and GTP poised for catalysis (IC3). (A) The bubble LSP in IC3. The DNA regions interacting with POLRMT, TFB2M and TFAM are indicated. The non-template, template, and RNA/NTP bases are in cyan, pink, and green, respectively. The hollow circles represent untraced nucleotides. The position of the active site (*) is indicated. (B) Left: Unsharpened experimental cryo-EM density map of IC3 at 5σ colored by protein and DNA chains – POLRMT (light blue), TFB2M (yellow), TFAM (brown), non-template DNA (cyan), and template DNA (pink). Right: ∼180° turn view of the unsharpened map; the POLRMT domains/motifs are colored as: pentatricopeptide repeat (PPR - pale green), intercalating hairpin (ICH – purple), specificity loop (magenta), fingers (blue), thumb (green), and palm (red). (C) Cartoon representation of IC3 structure. The DNA promoter in surface representation shows a fully resolved transcription bubble with 2-mer RNA and GTP (green) based paired with the template. (D) One-dimensional representation of POLRMT, TFB2M and TFAM with its domains and key structural elements colored. The untraced protein regions are in white.

The three component proteins POLRMT, TFAM, and TFB2M of the h-mt initiation complex were assembled on the bubble LSP promoter and purified by size-exclusion chromatography (SEC) (Figures S1D-S1F). The SEC-purified complex was incubated with GTP and 3’-dATP and supplemented with additional TFAM for cryo-EM structural study, as described in the experimental section. The resulting 3.4 Å resolution cryo-EM structure (Figures 2B and 2C) captured a fully resolved transcription bubble and a catalytically synthesized pppGp(3’-d)A RNA and an incoming GTP base-paired with the +1 to +3 template (Figures 2C, S2A-B). This initiation intermediate is referred to as the IC3.

The following protein and DNA regions were modeled into the experimental cryo-EM density of the IC3 structure; DNA from −38 to +12, TFB2M from residues 71 to 396, POLRMT from residues 122 to 1230 (except the disordered linker 152-158, linker loop 200-216, and tip of the thumb 741-754), and TFAM from amino acids 43 to 234 (except disordered regions 171-176 and 235-246) (Figure 2D).

The overall assembly of POLRMT, TFB2M, and TFAM proteins in IC3 is similar to that observed in the earlier crystal structures of the open complex^27^; TFAM binds to an upstream site on the LSP and bends the DNA into a hairpin structure while interacting with the POLRMT tether helix (residues 122-145) (Figures 2B and 2C, Figure S2A). The pentatricopeptide repeat (PPR) domain (residues 218-368) interacts with the upstream DNA region −16 to −11. Our structure resolved an additional region of POLRMT (159-199) between the tether helix and the PPR domain as two flexible helices (Figures S2C and S2D); this region was disordered in the previous crystal structure. POLRMT interacts with TFB2M through a lever loop or B2-hairpin (residues 588-604) analogous to the Mtf1-hairpin in y-mt ICs^24,26^, and this loop is now defined in the structure. The intercalating hairpin (residues 605-624) and specificity loop (residues 1085-1108) in POLRMT interact with the −7 to −3 region of the LSP and are responsible for promoter specificity (Figure 2A, Figures S2C and S2E). The structure also resolved a previously unidentified TFB2M region between β6 and β7 (residues 268 to 294) positioned near the downstream DNA (Figures S2C and S2F). The C-terminal polymerase domain of POLRMT was defined by the palm (residues 790-953 & 1108-1230), thumb (residues 705-790), and fingers (residues 937-1085) necessary for RNA synthesis (Figure 2D).

The local resolution of the TFAM-associated region in the IC3 structure was relatively low (∼4.5 Å) compared to the ∼3 Å resolution of the POLRMT/TFB2M core (Figure S3A). A total of 863 POLRMT Cα atoms in IC3 (excluding the fingers domain) were superimposed on the crystal structure (PDB: 6ERP) with an RMSD of ∼1 Å (Figures S3B), and this superposition revealed a ∼15 Å shift in the positions of TFAM between the cryo-EM and crystal structures (Figure S3C). However, superposition of the TFAM showed no noticeable differences in the TFAM interactions with the upstream DNA or to the tether helix (Figure S3D). Thus, the shift in TFAM positioning in relation to POLRMT is due to the flexible linkage with the tether helix. The DNA acts as a hinge, allowing TFAM to move as a rigid body relative to POLRMT.

The cryo-EM sample showed the co-existence of another IC3 population lacking density for TFAM and the tether helix. The population of the TFAM-containing complex was significantly reduced when the SEC-purified POLRMT/TFAM/TFB2M/DNA complex was not supplemented with additional TFAM during grid preparation. Importantly, in the population lacking TFAM, the upstream DNA was straight rather than bent suggesting the dissociation of TFAM (Figure S3E). The flexible tether-helix linkage and the DNA bending-unbending transitions of TFAM^33,34^ may have caused TFAM to dissociate from the complex. Nevertheless, IC3 structures with and without TFAM had a highly superimposable POLRMT/TFB2M/DNA core structure that aligned with an RMSD of 0.3 Å for 849 Cα-atom superposition. Henceforth, we discuss the TFAM-containing IC3 structure unless otherwise noted.

### POLRMT makes promoter specific interactions with the template strand

Positioning the template in the active-site cleft of IC3 involves bending of the DNA by ∼55° between the upstream and downstream DNA duplexes stabilized by interactions with POLRMT and TFB2M (Figure 3A). The upstream arm of the bent DNA, extending from the TFAM-anchored position to the transcription bubble, is stabilized by the PPR domain and the specificity loop of POLRMT, as well as interactions of TFB2M with the −7 and −8 base pairs (Figure S2E). The downstream DNA is stabilized by positively charged lysine and arginine residues (R1003, R1007, R1034, R1059, R1113, K1114, and K1116) from the fingers and palm domains of POLRMT and (R276, K277, K284, and R285) from the newly resolved TFB2M loop (residues 268 - 294) (Figure 3B).

**Figure 3.**
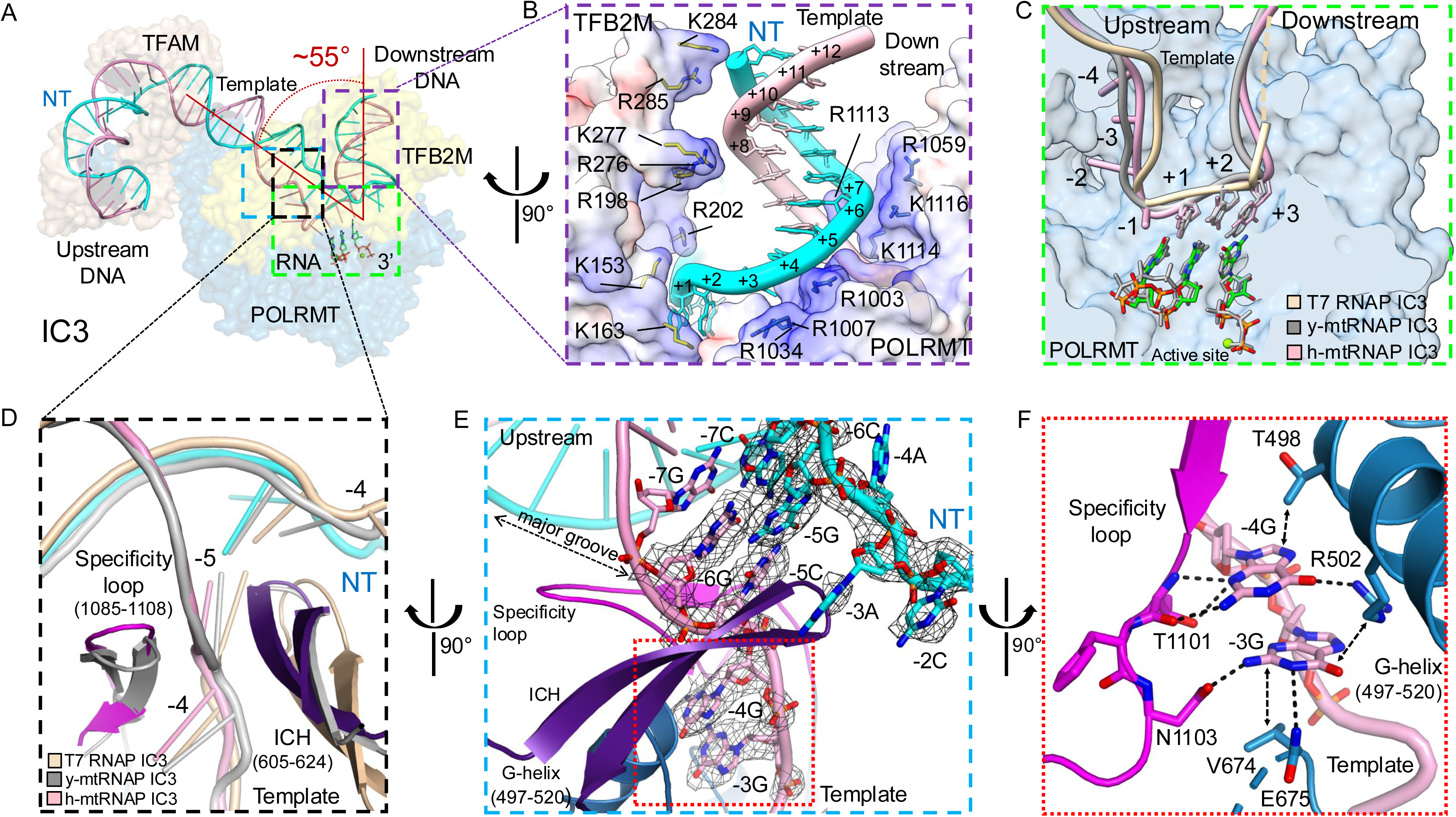
The IC3 defines the initiation bubble and the transcription start site. (A) DNA track (template – pink; non-template – cyan; RNA – colored by heteroatom, stick representation) in IC3 reveals ∼55° bend between the upstream and downstream DNA. Surrounding proteins shown as transparent surfaces (TFAM – brown; TFB2M – yellow; POLRMT –blue). The boxed regions are zoomed in subsequent panels. (B) Zoomed view showing the interactions of the downstream DNA. The electrostatic potential surfaces of POLRMT (right) and TFB2M (left) show an arginine/lysine rich positively charged groove accommodating the duplex DNA. (C) Zoomed view of the characteristic U-shaped template track base-paired with RNA from +1 to +3 shows a structurally conserved initiation bubble in the IC3 of h-mt (pink and green), y-mt (gray) (PDB:6YMW), and T7 (tan) (PDB:1QLN) aligned based on their RNAP Ca superpositions. (D) Zoomed view of the upstream edge of the initiation bubble in the IC3 of h-mt (colored), T7 (tan), y-mt (gray) aligned by RNAP Ca suggests −4 base-pair as a conserved upstream promoter melting position in single-subunit RNAPs. The intercalating hairpin (ICH – purple) wedges between the -5 and −4 positions of the bubble, while the specificity loop (magenta) stabilizes the upstream DNA. (E) The experimental density map (8σ) for the promoter region (-6 to −2) defines the upstream edge and the melted −3 and −4 bases in IC3. (F) Zoomed view highlighting the base-specific interactions of the template −4 and −3 guanines with the specificity loop and G-helix of POLRMT. Hydrogen bonds and hydrophobic interactions are indicated by dotted lines and arrows, respectively.

The template DNA positioned at the active site assumes a characteristic U-shaped track, conserved in POLRMT, y-mt Rpo41, and T7 RNAP chains in their respective IC3 states^23,24,26^ (Figure 3C). In all three systems, the transcription bubble begins with the melting of the −4 base pair, showing a structurally conserved initiation bubble in single-subunit RNA polymerases. The intercalating hairpin (ICH) of POLRMT stabilizes the upstream edge of the transcription bubble by wedging between the -5 base pair and the melted −4 nucleotides (Figure 3D).

The template DNA is stabilized in the IC3 through upstream base-specific interactions with the specificity loop, which is fully resolved. The specificity loop inserts into the major groove of the upstream DNA between the −7 and -5 base pairs making base-specific interactions with the - 4 and −3 guanines in the melted template strand (Figures 3D-3F and S2E). Biochemical studies showed these guanines are critical for transcription initiation (Figures 1B and 1C). The N2-amino group and N3 atom of the −4 guanine form H-bonds with the peptide backbone and the side chain of T1101 in the specificity loop (Figure 3F). Similarly, the N2-amino group of the −3 guanine H-bonds with the N1103 side chain. Furthermore, the −4 and −3 guanines are recognized by T498 and R502 of the POLRMT G-helix (residues 497-520) via H-bonds and hydrophobic interactions; the R502H mutation reduces transcription initiation efficiency^28^. The −3 guanine H-bonds with the E675 main chain and the stacked −4 and −3 guanine bases are flanked from both ends by the side chains of T498 and V674, respectively, providing additional stability (Figure 3F).

### IC3 structure reveals that transcription starts from the +1 position

The IC3 captured a *de novo* synthesized 2-nt RNA pppGp(3’-d)A and GTP base-paired to the +1 to +3 template positions (Figures 4A, 4B (left), 4C (left) and S2B). The added nucleotides GTP and 3’-dATP could initiate RNA synthesis from either the −1 or +1 position of the bubble LSP template sequence (-1)CCTCT. However, no RNA density was observed at the −1 position. This is consistent with the transcription assays showing negligible synthesis of pppGpG (from position - 1) relative to pppGp(3’-d)A (from position +1) (Figure S1B), reaffirming that −1 position is not a start site. The structure indicates that the anchored template −4G and −3G to POLRMT (Figure 3F) would not permit the −1 and +1 nucleotides to stretch and engage complementary NTPs at the active site for *de novo* RNA synthesis. Thus, the IC3 structure provides an atomic resolution snapshot of the transcription initiating from +1 TSS on LSP.

**Figure 4.**
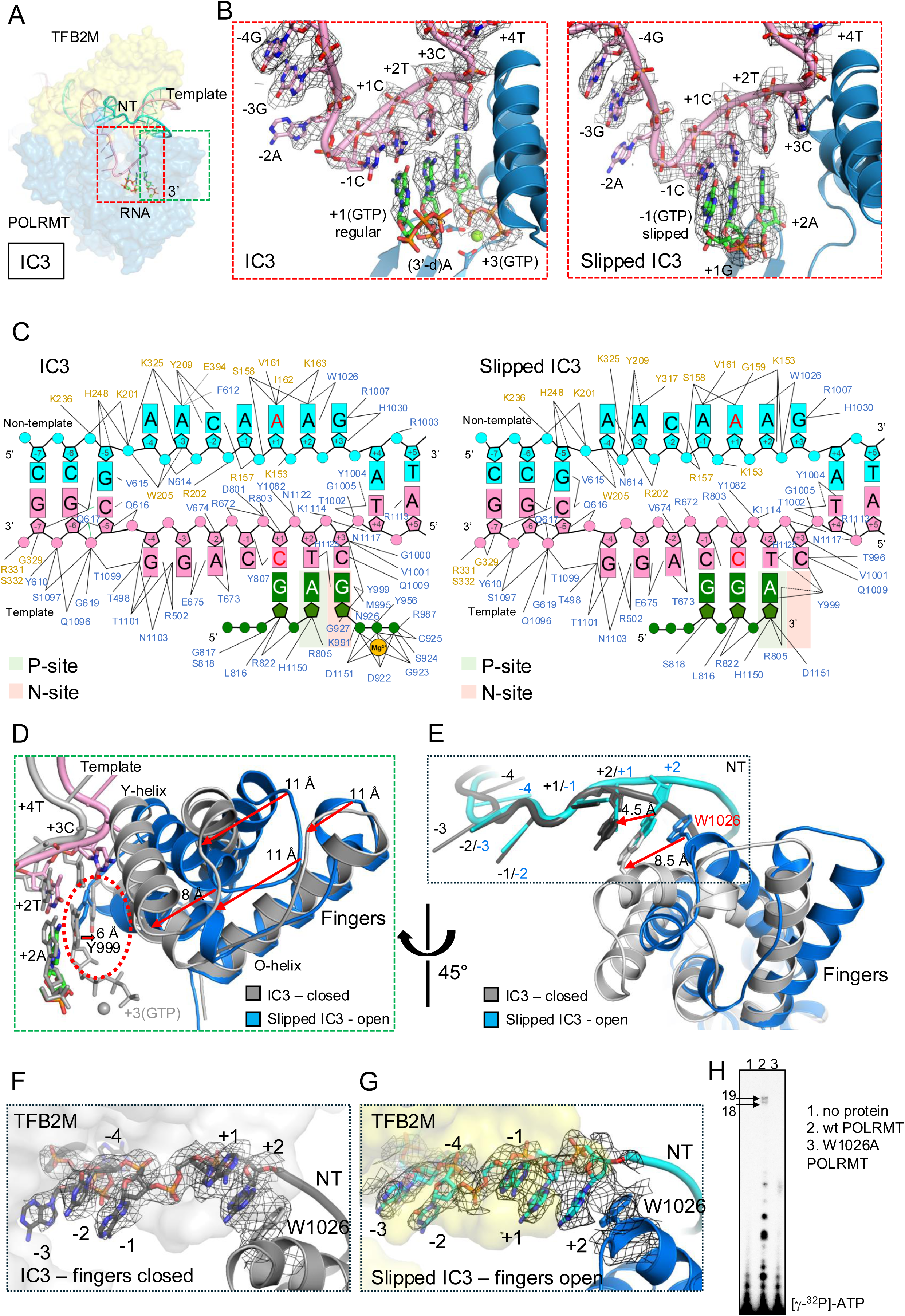
Transcription bubbles in IC3 and slipped IC3 reveal non-template repositioning with fingers opening and closing. (A) IC3 structure indicating the regions zoomed in subsequent panels; TFAM is omitted for clarity. (B) The cryo-EM density maps of IC3 at 6s (left) and slipped IC3 at 2.5s (right) define the RNA positions with respect to the unaltered template track. The RNA starts from +1 in IC3 and −1 in slipped IC3. The fingers Y helix (in blue) interacts with the template and GTP. The triphosphate of −1 GTP in RNA, pppGpGpA, is omitted for clarity (right). (C) Schematic representation of the transcription bubbles in IC3 (left) and slipped IC3 (right) and interacting protein residues from POLRMT (blue) and TFB2M (yellow); cyan non-template, pink template, green RNA. The P-site (priming site) and N-site (nucleotide-binding site) are shaded light green and red, respectively. (D) The superimposed IC3 (gray) and slipped IC3 (colored) aligned through POLRMT shows fingers-closed (gray) and fingers-open (blue) states in respective structures. In the slipped IC3, Y999 occupies the empty N-site. In IC3, GTP is bound at the N site and Y999 is repositioned with the closing of the fingers domain, as indicated by the red arrows. (E) A view at about 45° clockwise of panel D shows fingers opening and closing impacts the positioning of the non-template (NT) strand in the bubble. The residue W1026, at the tip of the fingers, stacks with NT +2A base. As the fingers close (blue è gray), W1026 moves ∼8.5 Å along with the stacked +2 nucleotide, causing a registry shift of the bases in the single-stranded NT region from −3 to +2. (F and G) Cryo-EM density maps at 6s (left) and 2s (right) define the NT −4 to +2 region and W1026 in fingers open and closed states. TFB2M interacts with the NT and is represented as gray and yellow molecular surfaces in fingers closed and open states, respectively. (H) Transcription runoff assay showing W1026A POLRMT is defective in abortive and run-off synthesis.

The *in situ* synthesized 2-nt RNA pppGp(3’-d)A lacks the reactive 3’-OH group; hence, the next nucleotide GTP was captured at the nucleotide-binding site (N-site), base-paired with the template +3 nucleotide in a state just before the phosphodiester bond formation. The fingers of POLRMT is in the closed conformation (Figure 4B (left), 4D). The GTP makes several contacts with the fingers domain O-helix (Figure 4C, left), which is critical for catalysis and conserved in single-subunit polymerases^35,36^. The triphosphate of GTP interacts with Y956, R987, and K991 of the fingers domain and chelates a Mg^2+^ ion (ion B), which is also coordinated with the catalytic residues D1151 and D922, and main-chain carboxyl of G923. The GTP base is stacked with Y999 residue of the O-helix in a conformation typical to the closed-fingers state (Figure 4C, left). This stacking of the incoming NTP is conserved in the elongation complex^37^. The Y999 is highly conserved in single-chain polymerases (Y639 in T7 RNAP and Y1022 in y-mtRNAP) and is critical for recognizing the 2’-OH group of an incoming NTP^23,26^.

### Slipped IC3 structure reaffirms the catalytic addition between +1 and +2 positions

To determine the structure of a slipped state, we added a 2-nt RNA pppGpG to base pair with the template −1C and +1C of the LSP bubble, and an incoming ATP to base pair with template +2T. Transcription assays indicated that pppGpG is elongated to a 3-nt RNA in the presence of ATP (Figure S1C). For this structure, during cryo-EM grids preparation, we did not supplement the sample with additional TFAM. Hence, we obtained a POLRMT/TFB2M/LSP bubble/pppGpGpA (3nt-RNA) slipped IC3 complex with no bound TFAM as the predominant population. We refer this as slipped complex here onwards.

The slipped structure, determined at 3.15 Å resolution, revealed that the 3-nt RNA produced from the reaction between pppGpG and ATP is base-paired with the template from −1 to +2 positions. As in IC3, the phosphodiester bond forms between the nucleoside at the +1 position and NTP at the +2 position, confirming that the first nucleotide addition starts from the +1 position. The 3’-end of the RNA is positioned at the P-site; thus, the 3-bp RNA:DNA in the slipped structure is captured in the post-translocated state (Figures 4B (right) and 4C (right)). Consequently, the +3 base-pair is melted, but the +3 template base has not reached the N-site to base-pair with an incoming NTP that was not provided. In the absence of an NTP at the N-site, the fingers domain in the slipped structure is captured in the open conformation (Figure 4D).

Although, the 5’-end of RNA is positioned at the −1 site in the slipped structure, the upstream edge of the bubble is still melted from the −4 position as in IC3 (Figure S2B (right)). The overall size of the transcription bubble, the template track, and its interactions with the proteins are similar to those in the IC3 structure (Figure 4B and 4C). Thus, the transcription bubble in slipped structure is not perturbed in its shape or interactions with proteins. As a result, RNA slippage does not require overcoming a significant energy barrier.

### Fingers opening and closing repositions the non-template strand

The IC3 and slipped structures captured the POLRMT fingers in closed and open states, respectively. As observed in all single subunit polymerases, the fingers domain positionally cycles between closed and open states at each nucleotide addition step^26,38^. The fingers domain closes to hold an NTP and opens when the N-site is empty following the RNA:DNA translocation. Thus, in the IC3 structure with GTP bound at the N-site, the fingers domain is closed, whereas in the slipped structure i.e. with an unoccupied N-site, the fingers domain assumes the open state (Figure 4D). In the fingers open conformation, the POLRMT residue Y999 at the base of the O-helix occupies the N-site. In the fingers-closed IC3 structure, Y999 moves out to accommodate an NTP in the N-site.

The tip of the fingers contains a residue W1026, which stacks with the non-template +2A in both the IC3 and slipped structures (Figures 4E-4G). When the fingers open and close, W1026 moves by about 8.5 Å, while continuing to stack with the non-template +2A (Figure 4E). The W1026 repositioning with fingers opening/closing results in a rearrangement of the non-template strand −4 and +1 nucleotides interactions with TFB2M, as discussed below. This non-template rearrangement is a surprising and unique finding in the h-mt IC and is not observed in y-mt ICs. Functionally, W1026 is an essential residue for transcription initiation (Figure 4H). This residue is conserved in vertebrate POLRMT but absent in T7 and y-mt RNAPs.

### TFB2M makes promoter specific interactions with the non-template strand

The single-stranded non-template nucleotides from −4 to +1 interact with TFB2M through contacts involving the nucleotide backbone, base-stacking, and hydrogen bonding (Figure 4C). Two structural elements of TFB2M are primarily responsible for these interactions: the NT- stabilizing loop (K_153_LDPRSGGVIKPP_165_), rich in proline and glycine, and the adjacent helix encompassing residues E200 to Y212 (Figure 5A-5D). The NT-stabilizing loop interacts with the non-template DNA sequence (-1)AAA(+2), conserved in all three h-mt promoters^7^. The proline residues guide the central flexible S_158_GGV_161_ region of the NT-stabilizing loop to the vicinity of the −1 to +2 non-template nucleotides. AlphaFold models^39^ indicates that the loop is structurally conserved in vertebrates (Figures S4A and S4B), suggesting that the general roles of the NT- stabilizing loop elucidated by the current structures are likely conserved in vertebrate mitochondrial ICs.

**Figure 5.**
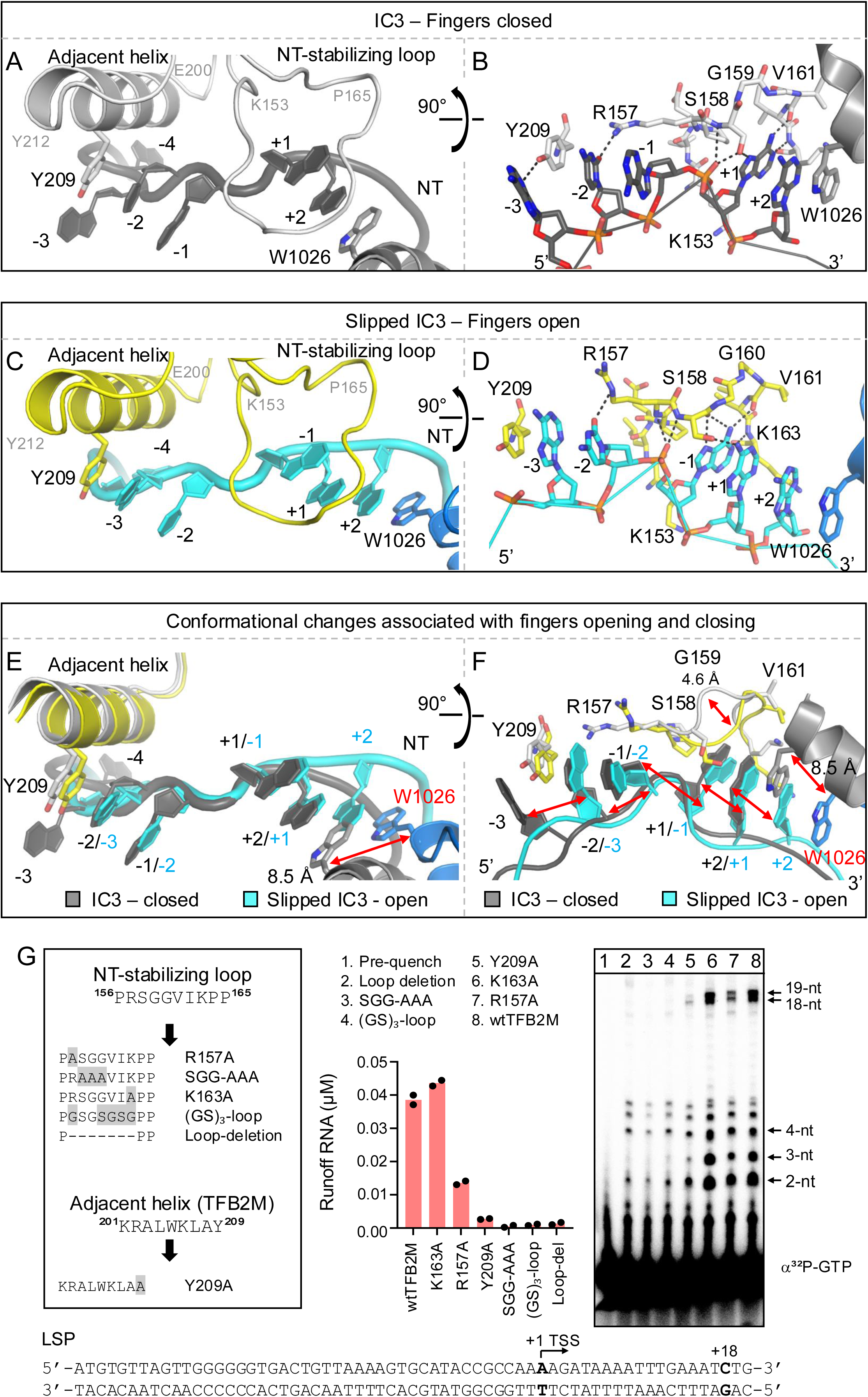
Structural elements of TFB2M interacting with the non-template −3 to +1 nucleotides are important for transcription initiation. (A) The NT-stabilizing loop (153-165) of TFB2M interacts with the non-template +1 and +2 adenosines (dark gray) in IC3. POLRMT W1026 stacks with +2A. The Y209 in the adjacent helix (200-212) of TFB2M stacks with −2C base. (B) Atomic model shows specific interactions between TFB2M and non-template in IC3; the dotted lines denote H-bonds. (C) The NT-stabilizing loop of TFB2M (yellow) interacts with the non-template −1 and +1 adenosines (cyan) in slipped IC3. The Y209 in the adjacent helix now stacks with the −3A base. (D) Atomic model shows the extensive base-specific interactions of the NT-stabilizing loop of TFB2M (yellow) with −1A of the non-template (cyan) in slipped IC3; the dotted lines denote H-bonds. (E and F) Alignment of IC3 and slipped IC3 based on POLRMT Ca superposition shows the repositioning of non-template guided by fingers open and closed states. Fingers closing upon NTP binding, shifts the W1026/+2A entity that consequently pushes the non-template −4 to +1 nucleotides to slide over the TFB2M NT-stabilizing loop and Y209, and redefines the −3 to +1 nucleotide interactions. The shifts are indicated by red arrows. (G) Validation of the key residues in the TFB2M structural elements interacting with the non-template by transcription runoff assays.

The non-template strand interactions with TFB2M have rearranged between the fingers closed and open states, observed in IC3 and slipped structures, respectively (Video S1). The W1026 movement with fingers opening and closing causes a registry shift for each non-template base from positions −3 to +2. As a result, the −3 to +1 base positions in the fingers-open align with the −2 to +2 base positions in the fingers-closed structure (Figure 5E and 5F). Additionally, the NT- stabilizing loop interacts with the +1A in the finger-closed IC3 state (Figure 5A) and with the −1A in the finger-open state (Figure 5C).

In the fingers-closed IC3 state (Figure 5A and 5B) (i) W1026 side-chain, non-template +2A, and +1A are stacked, (ii) the +1A has base-specific H-bond interactions with the NT-stabilizing loop, (iii) the non-template −2 base stack with the −1 base and the aromatic-ring side-chain of Y209 from the TFB2M adjacent helix, and (iv) −2 and −3 bases form H-bonds with the R157 and Y209 side chains of TFB2M, respectively. The −4 base lacks significant interactions and thereby is weakly defined.

In the fingers-open state (Figure 5C and 5D) (i) W1026 side chain, non-template +2A, +1A, and −1A are stacked, (ii) The non-template −3 base stacks with the Y209 side-chain, and (iii) the - 1A at the center interacts extensively and base-specifically with the flexible S_158_GGV_161_ tip of the NT-stabilizing loop. The N6 amino group and N1 atoms of −1A form base-pair like H-bonds with main-chain carbonyl and amino groups of V161 and K163, respectively (Figure 5D). The N6 and N7 of −1A also interact with the main-chain carboxyl of S158 and one of the −1A phosphate oxygen forms H-bonds with the main-chain amino groups of R157 and S158. The Oγ atom of the S158 interacts with the N6-amino group of the +1 adenine.

### Interactions of the NT-stabilizing loop and the adjacent helix are important for IC formation

It is reasonable to assume that at the promoter melting stage when no NTP is bound, the fingers domain is in the open state. Thus, in the open complex, the NT-stabilizing loop is expected to interact base-specifically with the non-template −1 adenine. Our biochemical assays indicate that non-template −1A is essential and its substitution by T/G/C or abasic nucleotide disfavors transcription initiation from +1 TSS (Figures 1C and 1E). However, substitution of +1A to other bases is well tolerated; but deletion of the base eliminates slippage synthesis (Figure 1E). Thus, base-specific interactions of the non-template −1A with the NT-stabilizing loop are critical for promoter melting and for the correct positioning of the template +1 and +2 nucleotides for NTP binding and transcription initiation. In the absence of −1A interactions, the NT-stabilizing loop is likely engaged with the +1 or +2 base, causing a downstream shift of the TSS to yield shorter 17- and 16-nt RNA transcripts, respectively, as observed in our transcription runoff assay (Figures 1C and 1E).

The functional significance of the NT-stabilizing loop and the adjacent helix residue Y209 were assessed through mutagenesis and transcription runoff analyses (Figure 5G). The NT- stabilizing loop was mutated in several ways: (i) a loop deletion mutant was created by removing the residues between the two prolines (P_156_RSGGVIKP_164_), (ii) the loop was replaced with a (GS)_3_- loop i.e., G_157_SGSGSG_163_ that preserved the S158 interactions but eliminated those of R157 and K163, (iii) the SGG sequence was replaced with AAA to reduce loop flexibility and disrupt S158 interactions with the −1 and +1 non-template adenine bases, and (iv) TFB2M point mutations, R157A, K163A, and Y209A, were introduced. Each TFB2M mutant was purified to homogeneity and tested in transcription runoff assay on the duplex LSP with POLRMT and TFAM.

The (i) to (iii) mutations in the NT-stabilizing loop abrogated runoff products (lanes 2 – 4, Figure 5G), indicating the essential role of the loop interactions in transcription initiation. A low amount of shorter RNA products (4- to 6-mers) produced by these mutants could be from non-specific initiation. The point mutation K163A showed no defect in transcription initiation and produced similar amounts of 18-nt and 19-nt runoff products as the WT TFB2M (Figure 5G). The R157A and Y209A mutants showed 35% and 7% of runoff products relative to WT TFB2M, respectively. These results confirm that the interactions of the NT-stabilizing loop and the residue Y209 identified in the structures are crucial for productive transcription initiation.

### TFB2M mutants are defective in promoter melting and transcription initiation

The reduced transcriptional activity observed in TFB2M mutants may be caused by a failure in assembling the TFB2M/POLRMT/DNA/TFAM complex or defects in promoter melting or the correct positioning of the non-template strand in the active site for initiation. We developed a fluorescence polarization assay to quantify the binding affinity of TFB2M to the POLRMT/TFAM/LSP complex^19,40^ (Figure 6A). TFB2M was labeled with Alexa-488 fluorophore and its binding affinity to POLRMT was measured by fluorescence polarization after adding increasing amount of POLRMT/TFAM/LSP complex. The titration experiment indicated that WT TFB2M binds to the complex with a K_D_ of 6.1 nM. Subsequently, the binding affinities of the mutant TFB2Ms were measured through competition assays (Figure 6B). Here the complex of labeled WT TFB2M/POLRMT/TFAM/LSP was added to increasing concentration of individual unlabeled mutant TFB2M. All the mutants, including defective Y209A, and transcriptionally inactive SGG- AAA, and (GS)_3_-loop mutants competed effectively with the WT TFB2M, providing inhibition constant (K_i_) values ranging from 25 nM to 64 nM, within the range of the WT TFB2M (Figure 6B). Based on the results, we conclude the reduced or lost transcription activity of TFB2M mutants is not caused by a failure to form the complex of POLRMT/DNA/TFAM with a mutant TFB2M.

**Figure 6.**
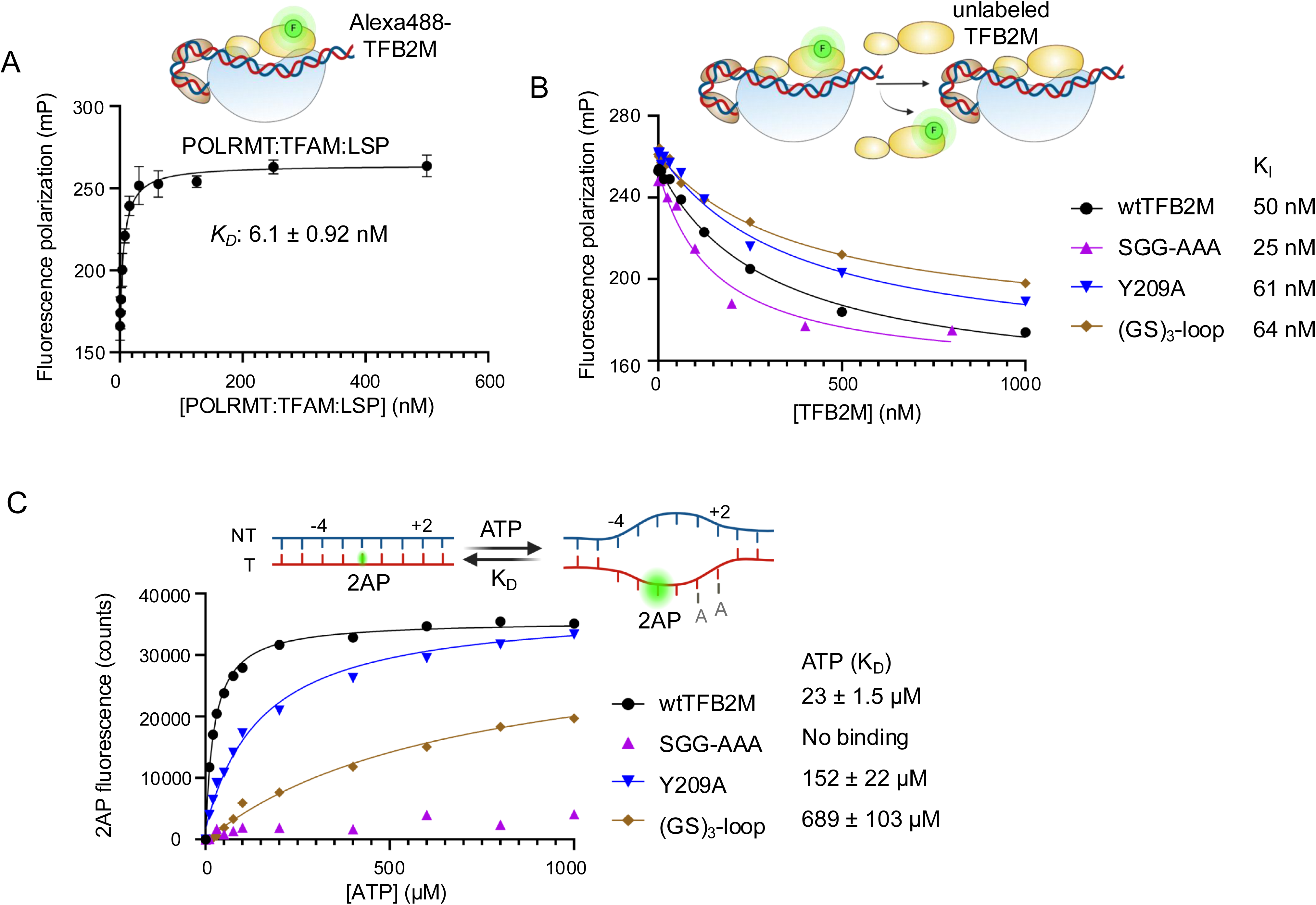
The NT-stabilizing loop and the adjacent helix are essential for promoter melting and initiation complex formation. (A) Quantitation of TFB2M binding affinity to LSP/POLRMT/TFAM by fluorescence polarization. (B) Quantitation of TFB2M mutant binding affinity to LSP/POLRMT/TFAM complex by a competition assay. The competition provided the IC_50_ which were used to calculate the K_I_ values as outlined in the experimental methods. (C) Quantitation of initiating ATP binding affinity to 2-aminopurine (2AP) modified LSP. The 2AP fluorescence increases upon promoter melting and ATP binding and provides the ATP K_D_ values. The experiments were conducted in triplicate or duplicate.

Next, we employed the fluorescence-based 2-aminopurine (2AP) promoter melting assay to assess whether the above TFB2M mutants are defective in forming an initiating complex (Figure 6C). Previous studies have shown that promoter melting and initiating ATP binding cause a significant increase in the fluorescence intensity of 2AP at LSP −2 position (previously annotated as −1)^19^. The initiating ATPs bind at the polymerase active site only when they base pair with the melted +1 and +2 template bases. Any defect in promoter melting or template alignment would decrease the binding affinity of the initiating ATPs. The −2 C:2AP LSP was incubated with POLRMT, TFAM, WT (or mutant) TFB2M, and added to increasing concentrations of ATP (Figure 6C). The WT TFB2M complex showed high-affinity ATP binding with a K_D_ of 23 µM. In contrast, the Y209A mutant complex showed a ∼7-fold weaker ATP binding affinity (K_D_ of 152 µM), the (GS)_3_-loop mutant complex showed a ∼30-fold weaker affinity (K_D_ of 689 µM), and the SGG-AAA mutant complex showed no ATP binding, even at concentrations up to 1 mM. These results are consistent with the structural findings and indicate that the base-stacking interactions of Y209 with the non-template −2 or −3 bases and the NT-stabilizing loop interactions with the −1 to +2 non-template adenines are essential for promoter melting and consequently for initiating complex formation.

Introducing a mismatch at −2, +1, or +2 position partly rescued the runoff transcription activity of R157A, Y209A, and the (GS)_3_-loop mutant TFB2Ms, which showed low or undetectable activity on the duplexed LSP (Figure S4C). However, none of the mismatches could rescue the activity of the SGG→ AAA mutation that eliminated the −1A base-specific interactions (Figure 5D). These results confirm the role of the TFB2M residues in promoter melting and highlight the significance of the S_158_GG moiety’s interaction with the non-template −1A to trap the melted non-template strand for efficient initiation. The structural flexibility of the NT-stabilizing loop is critical for its interactions with the non-template bases.

### Multiple states of POLRMT/TFB2M/TFAM complexes on LSP co-exist in solution

The cryo-EM single-particle analysis revealed additional complexes of POLRMT despite the homogeneity of the SEC-purified complex of POLRMT/TFAM/TFB2M on the LSP bubble (Figure S1E) and supplementing the complex with additional TFAM. The predominant states that were discerned included: State-I - POLRMT/TFAM/TFB2M/DNA/RNA/GTP (IC3); State-II - POLRMT/TFB2M/DNA/RNA/GTP (IC3 – no TFAM); State-III - a homodimer of State-II interacting via the POLRMT PPR domain; State-IV a heterodimer of State-II with a POLRMT dimerized at their PPR domains; and State-V – free POLRMT monomers; with the cryo-EM maps calculated at 3.4, 3.5, 4.2, 4.2, and 3.6 Å resolution, respectively (Figures S5). More than 50% of the particles captured the IC3 with or without TFAM state (States I or II), ∼27% of the particles were of free POLRMT (State-V), and the remaining ∼18% represented two dimeric states (States III and IV). The existence of multiple states indicates that POLRMT in the initiation complex POLRMT/TFAM/TFB2M/DNA/RNA/GTP is in a dynamic equilibrium with several other functional states including the apo POLRMT and PPR-domain-interfaced dimeric POLRMT (Figure S6A). A recent study reported POLRMT dimers bound to the non-coding 7S RNA, which are inactive in RNA synthesis^41^. The 7S RNA-mediated dimers involved a different interface (ICH, specificity loop, and thumb) (Figure S6B). An analogous phenomenon is observed for cellular RNAP Pol I involved in ribosomal RNA synthesis^42^, where monomeric and dimeric states regulate its activity; the inactive dimeric state is thought as a storage pool converted to the active monomeric state in response to physiological changes. It is reasonable to assume that the dimeric states of POLRMT would reduce the availability of free POLRMT to participate in IC formation. Thereby, the relative heterogeneous distribution of monomeric vs. dimeric POLRMT states generated from purified homogenous h-mt IC in our cryo-EM sample may suggest that the POLRMT concentration and oligomerization have functional implications in the formation of the transcription initiation complex and the transcription initiation process.

## DISCUSSION

We have captured the structures of two critical intermediates, IC3 and a slipped IC3, in the human mitochondrial DNA transcription initiation pathway (Figure 7). These intermediates visualize the transcription bubble and provide insights into the mechanism of promoter-specific transcription initiation by POLRMT and TFB2M. This study provides the structural basis for two modes of initiation on the LSP sequence – the *de novo* RNA synthesis from the +1 position (nucleotide 406) and slippage synthesis. Fully resolved transcription bubbles, observed here for the first time in an h-mt POLRMT initiation complex, reveal that LSP is melted starting from the −4 position, similar to that in T7 and y-mt ICs, establishing now a conserved initiation bubble structure across single-subunit RNA polymerases. The promoter DNA is melted cooperatively by POLRMT and TFB2M through base-specific and base-stacking interactions with the −4 to +2 region of template and non-template. Specifically, POLRMT interacts with the −4 and −3 guanines in the template strand and +2 adenine in the non-template, while the NT-stabilizing loop of TFB2M is engaged with the non-template −1 adenine, aiding promoter melting and the alignment of the +1 and +2 template positions with the two initiating NTP at the polymerase active site to initiate *de novo* RNA synthesis from the +1 TSS (Figure 7, A→C). Our structures suggest that the −1 and +1 template are unlikely to engage NTPs at the polymerase active site to initiate *de novo* RNA synthesis from the −1 position. The interactions of TFB2M NT-stabilizing loop and POLRMT W1026 with the non-template (-1)AAA(+2) sequence, which is conserved in all three h-mt promoters (LSP, HSP, and LSP2), regulates RNA synthesis from +1 TSS.

**Figure 7.**
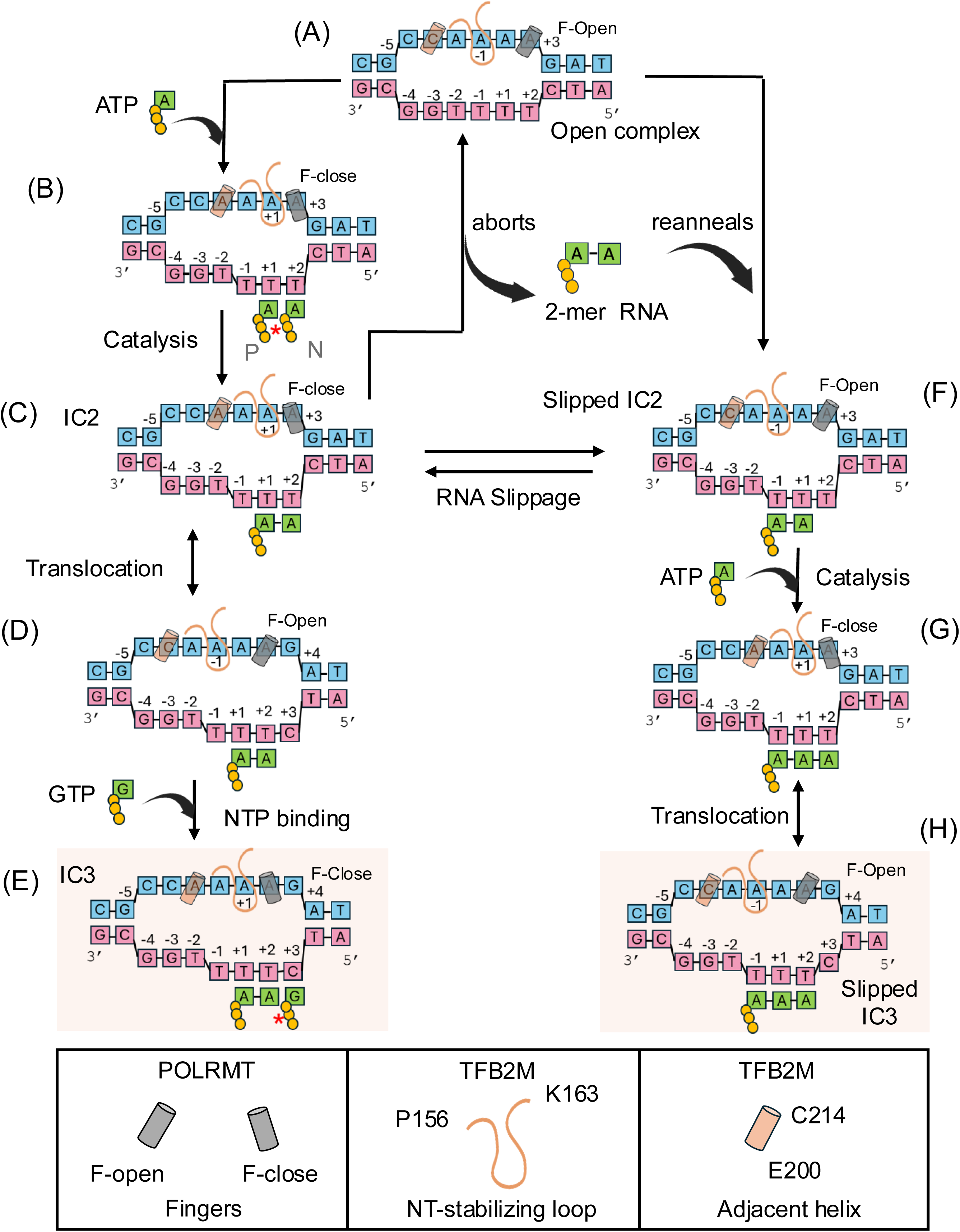
Transcription initiation mechanism. The initiating nucleotide (ATP) binds to the LSP open complex at the +1 priming site (P-site) and the +2 nucleotide-binding site (N-site), forming an initiation complex (A è B). This catalyzes the formation of IC2 (B è C), with a 2-nt pppApA RNA bound to the +1/+2 template positions. The RNA in IC2 can abort (Figure S1B), slip, or elongate into a 3-nt RNA. The IC2 is elongated to the IC3 observed in this study following the sequence of events of downstream DNA melting, translocation, and nucleotide binding (C è D è E); the catalytic incorporation was blocked by the 3’-deoxy end of the RNA in the IC3 structure. Alternatively, for slippage, the 5’-end of the 2- nt RNA repositions from the +1/+2 to base pair with the template −1/+1 (C è F), forming a slipped IC2 intermediate, which can then get elongated to slipped IC3 by incorporation of the next nucleotide requiring no additional DNA melting (F è G è H). If aborted, the 2-nt RNA reanneals with the template in the open complex to form IC2 or slipped IC2 intermediates that are interchangeable. As nucleotides bind to the N-site, the fingers move from open to close state, that consequently, shifts the TFB2M NT-stabilizing loop interactions from the non-template −1A to +1A. The right or left slanting of the gray helix represents the fingers’ open or closed states, respectively. The NT-stabilizing loop and adjacent TFB2M helix are denoted by an orange loop and helix, respectively, with orange shaded boxes highlighting the structurally obtained states.

Following *de novo* RNA synthesis from the +1 TSS, the 2-nt RNA in the IC2 intermediate has several possible fates (Figure 7). The 2-nt RNA may dissociate into solution as an abortive product, regenerating the open complex (C→A). The 2-nt RNA can be elongated to an IC3 intermediate following the +3 base pair melting and NTP binding (C→E), as captured in our study. Alternatively, the stretch of (-1)TTT(+2) in the promoter templates permits the 2-nt RNA to slip, repositioning its 5’-end at the −1 position (C→F). A slipped IC2 can also form through the reannealing of the aborted 2-nt RNA (A→F). The slipped IC2 binds the next NTP without downstream DNA melting (F→G). Thus, 3-nt RNA synthesis from slippage is expected to be energetically favorable than the 3-nt RNA synthesis without slippage, which requires downstream DNA melting. Accordingly, RNA-seq results show that the slippage product is the predominant transcript from all three promoters^7,28,29^.

The y-mt and h-mt RNAPs rely on structurally homologous transcription factors, Mtf1 and TFB2M, respectively, for promoter melting and stabilizing the initiation complex (Figure S6C). In general, both factors assist the promoter melting process by trapping the non-template strand of the initiation bubble using base-specific interactions that also guide site-specific transcription initiation^24,26,27^. However, there are notable differences in the promoter melting and non-template strand stabilization mechanisms between the two systems. The y-mt Mtf1 interacts with non-template −4 to −1 nucleotides (Figure S6C) by sandwiching the −2 guanine base between Y103 and W105 side chains and forming a network of base-specific H-bonds. The interactions of Mtf1 with the non-template remain stable throughout the transcription initiation process, from the opening of the bubble to the synthesis of an 8-mer RNA^24,26^. In contrast, TFB2M contains an NT-stabilizing loop and an adjacent helix, both absent in Mtf1, that interact with the non-template nucleotides. The NT-stabilizing loop of TFB2M protrudes over the non-template −1A by making base-specific interactions that are important for promoter melting and transcription start site selection. Following bubble opening, the interactions between TFB2M and the non-template strand are dynamic, linked with POLRMT fingers opening and closing movements. The stacking of W1026, at the tip of the fingers domain, with the +2 non-template base is responsible for repositioning the non-template strand and switching the −1/+1 non-template base interactions with the NT-stabilizing loop (Video S1).

The interactions of the NT-stabilizing loop with the −1/+1 non-template base are required for slippage synthesis, as evidenced by promoter mutational studies (Figure 1). The repositioning of the −1/+1 non-template bases over the NT-stabilizing loop is also like a slippage motion that is apparently coordinated with the RNA slippage at the template. It remains to be determined (i) if the NT-stabilizing loop shifts its interactions with the non-template to downstream nucleotides as the bubble expands in the later phase of initiation and help non-template scrunching, and (ii) if the dynamic interactions with the −1/+1 play a role in the transition from initiation to elongation phase. The W1026 also stacks with a non-template base in the elongation phase^37,43^. During elongation, the ratcheting motion of W1026 caused by the opening and closing of the fingers domain may be driving the coordinated melting of downstream DNA and reannealing of upstream DNA, maintaining the constant length of the RNA:DNA hybrid in the elongating bubble.

## LIMITATIONS OF THE STUDY

Mutation of the −4 non-template base decreases transcription efficiency (Figure 1); however, our structures do not reveal any base-specific interactions of the −4A with the proteins. This may be due to the −4C to −4A mutation in our study to create the bubble starting from the −4 position. It has been shown that POLRMT undergoes multiple cycles of slippage on the HSP, resulting in the addition of a run of adenines at the 5’-end of the HSP transcript whereas slippage is limited to only one nucleotide on LSP^19,28,32^. Our structures do not provide direct experimental evidence beyond single nucleotide RNA slippage event. The differences in the promoter sequence between LSP and HSP upstream of the −1 position likely account for these variations, which needs to investigated with further studies.

### Resource availability

#### Lead contact

Further information and requests for resources and reagents should be directed to and will be fulfilled by the lead contact, Smita S Patel and Kalyan Das.

#### Materials availability

Unique reagents generated in this study are available from the lead contact upon request with completed material transfer agreements.

#### Data and code availability

- Atomic models have been deposited in Protein Data Bank (PDB) and Electron Microscopy Data Bank (EMDB). Accession numbers are listed in Table S1 and in the key resources table. Raw gel image files and data are deposited in Mendeley data bank (link available).
- This paper does not report original code.
- Any additional information required to reanalyze the data reported in this paper is available from the lead contacts upon request.

## Acknowledgments

Funding for this work was provided by the National Institute of General Medical Sciences (NIGMS) [GM118086] to S.S.P., Canada Excellence Research Chairs (CERC) program to K.D., and Research Foundation – Flanders (FWO-Vlaanderen) doctoral fellowship 1162823N to Q.G.

## Author contributions

Conceptualization: S.S.P & K.D. Methodology: J.S. and Y.A., protein purification and assay development; Q.G., protein purification (for structural studies) and grid optimization; B.D.W., grid optimization, grid preparation and data collections. Investigation: J.S. and Y.A., biochemical assays; Q.G. & K.D., cryo-EM data processing, model building and structure analysis; S.S.P., biochemical assays and structure analysis. Visualization: all authors. Funding acquisition: S.S.P & K.D. Project administration and supervision: S.S.P & K.D. Original draft written by: J.S., K.D., & S.S.P, and later reviewed and editing by all authors.

## Supplemental Information

Document S1. Figures S1–S7

Table S1. Single Particle Cryo-EM data collection, refinement, and validation statistics Table S2. Oligonucleotide sequences

Video S1. Structural reorganization in human mitochondrial initiation complex associated with fingers opening and closing, related to Figures 1-5.

## STAR METHODS

### Proteins

Human POLRMT (43–1230, codon optimized), W1026A POLRMT mutant, and TFAM (43–245) were cloned into pPROEXHTb; TFB2M (21–396), Δ41 TFB2M (42–396), Δ59 TFB2M (60-396), and all NT-stabilizing loop and the adjacent helix TFB2M mutants were cloned into the bacterial expression vector pT7TEV-HMBP4 (kindly provided by Professor Miguel Garcia-Diaz at Stony Brook University School of Medicine).

The pPROEXHTB plasmid encoding N-terminal His6-tagged Δ42 POLRMT and W1026A POLRMT was transformed into *E. coli* BL21 Codon plus (RIL) competent cells (Agilent Technologies); and expressed and purified as previously reported^19,44^. Cells were grown at 37°C in LB (Luria-Bertani) liquid media broth, containing 100 μg/ml ampicillin, until an optical density (OD) at 600 nm reached 0.6–0.8 before induction with 0.3 mM isopropyl β-D-1-thiogalactopyranoside (IPTG, EMD Millipore Corp.). After an overnight incubation period at 18°C, cells were collected, resuspended in lysis buffer (40 mM Tris-HCl pH 7.9, 300 mM NaCl, 15% glycerol, 0.1% Tween-20, 1 mM phenylmethylsulfonyl fluoride (PMSF), 1 mM EDTA, 0.05 mM DDT) and lysed by sonification in the presence of protease inhibitors (1 tablet per 50 mL – Roche) and 1 mg/ml lysozyme at 30 sec ON/30 sec OFF pulses for 3 minutes and 4 repetitions. The resulting lysate was treated with 0.3% polyethyleneimine (PEI) to remove nucleic acids and other contaminants, followed by centrifugation. Subsequently, the supernatant was subjected to ammonium sulfate precipitation (55%) followed by centrifugation to pellet the precipitated proteins.

The ammonium sulfate pellet was resuspended in washing buffer (40 mM Tris-HCl, pH 7.9, 300 mM NaCl, 15% glycerol, 0.1% Tween-20, 0.3 mM EDTA, 0.05 mM DTT, 20 mM imidazole), filtered, and loaded on a 5 ml HiTrap DEAE FF and 5 ml HisTrap HP prepacked column (Cytiva) connected in tandem. After sample application the DEAE column was removed and the HisTrap HP column was washed with washing buffer. POLRMT was eluted with a gradient from 20 mM to 500 mM imidazole. Fractions containing POLRMT were identified by SDS PAGE and combined. The combined fractions were diluted to a NaCl concentration of 150 mM and loaded on a 5 ml Hitrap Heparin HP prepacked column (Cytiva). After washing the column with heparin washing buffer (40 mM Tris–HCl pH 7.9, 150 mM NaCl, 15% glycerol, 0.1% Tween-20, 1 mM PMSF, 1 mM EDTA, 1 mM DTT), POLRMT was eluted with a NaCl gradient from 150 mM to 1 M. The POLRMT eluent was collected and concentrated using a 30 kDa cutoff Amicon Ultra centrifugal filter (Millipore) and stored at −80°C.

TFAM was expressed and purified as previously reported ^19,44^. *E. coli* BL21 codon plus (RIL) cells transformed with the TFAM expressing vector were grown at 37°C in LB media, containing 100 μg/ml ampicillin, until O.D. at 600 nm reached 0.6-0.8 prior to induction with 1mM IPTG at 16°C for 16 hours. Subsequently, the cells were lysed in washing buffer (50 mM Na-phosphate pH 7.8, 300 mM NaCl, 10% glycerol, 20 mM imidazole, with or without 0.1% Tween-20) containing lysozyme (1 mg/ml) and protease inhibitor (1 tablet per 50 ml) by sonification (30sec ON/30 sec OFF for 4 repetitions). The clarified lysate was loaded on a HisTrap HP prepacked column (Cytiva), washed, and eluted with a 20 mM to 500 mM imidazole gradient. Fractions containing TFAM were collected and treated with a 1:100 or 1:30 (w:w) ratio of His-tagged TEV protease to TFAM for an overnight period at 4 °C. Afterwards, the sample was loaded on a 5 ml HisTrap HP prepacked column (Cytiva) and washed with washing buffer which eluted the cleaved TFAM. The eluent was diluted to a final concentration of NaCl below 150 mM, and subsequently loaded on a 5 ml HiTrap Heparin HP prepacked column (Cytiva), washed with heparin wash buffer (50 mM Na-phosphate pH 7.8, 200 mM NaCl, 10% glycerol) and eluted with a gradient of 200 mM to 1 M NaCl. The fractions containing pure TFAM were concentrated using a 10-kDa-cutoff Amicon ultra- centrifugal filter (Merck Millipore) and stored at −80°C.

The pT7TEV-HMBP4 plasmid encoding Δ20 TFB2M or its mutants (R157A, K163A, Y209A, loop-deletion (Δ157-163), and (GS)_3_-loop (R157G, G160S, V161G, I162S, K163G)), Δ59 TFB2M or its mutant (SGG-AAA (S158A, G159A, G160A)) and Δ41 TFB2M were transformed into *E. coli* strain ArcticExpress (DE3) cells (Agilent Technologie) and expressed and purified using the same protocol as previously reported^19,44,45^. The cells were grown overnight at 37°C in LB media with either 50 μg/ml kanamycin alone or a cocktail of antibiotics (50 μg/ml kanamycin, 20 μg/ml gentamycin, and 10 μg/ml tetracycline). The cells were then diluted 1:50 in fresh 1L LB media and grown without antibiotics for the kanamycin-only batch, or with the cocktail of antibiotics for the other batch, at 27°C until the OD reached 0.7–0.8. Induction was carried out by supplementing with 0.2 mM IPTG and growing the cells overnight at 14°C. Afterwards, cells were harvested by centrifugation and resuspended in TFB2M lysis buffer (20 mM HEPES–KOH pH 8, 1 M KCl, 5% glycerol, 10 mM imidazole) containing 1 tablet of protease inhibitor per 50 ml and 1 mg/ml lysozyme. After stirring for 1 hour, cells were lysed by sonication (30 sec ON/30 sec OFF for 3 minutes repeated 4 times), loaded onto a HisTrap HP prepacked column (Cytiva), washed with washing buffer (20 mM HEPES–KOH pH 8, 300 mM KCl, 5% glycerol, 20 mM imidazole), and eluted with a gradient from 20 mM to 500 mM imidazole. Fractions containing TFB2M were collected and treated with a 1:100 or 1:25 (w:w) ratio of His-tagged TEV protease to TFB2M for an overnight period at 4 °C. The sample was diluted to lower the KCl concentration below 200 mM and loaded on a HiTrap Heparin HP prepacked column (Cytiva). After loading, the column was washed (20 mM HEPES–KOH pH 8, 200 mM KCl, 5% glycerol, 1 mM EDTA, 1mM DTT), and eluted with a gradient of 200 mM to 1 M imidazole. The fractions containing TFB2M were combined, diluted to reduce the KCl concentration (< 200 mM), and loaded on a HiTrap SP HP prepacked column (Cytiva), washed with the same buffer as above, and eluted with a gradient of 200 mM to 1 M KCl. The fractions containing pure TFB2M protein were combined, concentrated using a 10-kDa-cuto off Amicon ultra-centrifugal filter (Merck Millipore) and stored at −80°C.

The molar concentrations of the proteins were determined using absorbance measurements at 280 nm in their respective buffers with the Nanodrop One UV-VIS spectrophotometer, utilizing the respective molar extinction coefficients of the proteins, and finally verified using SDS-PAGE gel (Figure S6D).

### In vitro runoff RNA transcription assay

Transcription assays were carried out at 25°C for 15 min with 1 µM POLRMT, 1.5 µM TFB2M or TFB2M mutants, 1 µM TFAM, and 1 µM LSP DNA template (-43 to +20) in reaction buffer containing 50 mM Tris acetate (pH 7.5), 50 mM sodium glutamate, 10 mM magnesium acetate, 1 mM DTT, 0.5 mM TCEP in the presence of an NTP mixture (ATP:100 µM, GTP:100 µM, UTP:100 µM, 3’-dCTP:400 µM and spiked with either [γ-^32^P]ATP (Figure 1B-left panel), or [α-^32^P]ATP (Figure 1B-right, C-left-middle panel & D panel) or [α-^32^P]GTP) (Figure 1C-right panel , E panel & S4D). GpG RNA (5 µM) was labeled at the 5’ end with [^32^Pi] with 2 µL of T4 Polynucleotide Kinase (PNK), 5 µL of 10X PNK buffer, and 4 µL of [γ-^32^P]ATP in a total volume of 50 µL. All the reactions were terminated by adding 125 mM EDTA and formamide dye solution (98% formamide, 0.025% bromophenol blue, 10 mM EDTA). Afterwards, the reaction mixtures were preheated at 95°C for 3 min and loaded onto 24% PAGE/4M urea gels. Subsequently, gels were exposed to a phosphor screen and visualized using the Typhoon FLA 9500 Phosphor-Image scanner (GE health care). The pre-quench reaction contains NTPs only. The RNA bands were quantified using ImageQuant software, and the amount of RNA product synthesized was determined using equation 1. All the experiments were performed in duplicates.

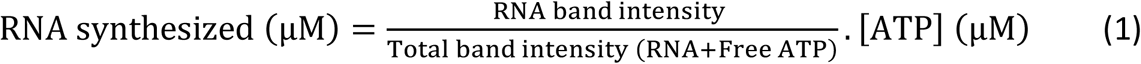

### ATP binding assay using 2AP fluorescence

A −2 C:2AP LSP (-43 to +20) modified DNA sample (D) (100 nM) was added to a premixed solution containing 150 nM of each of the following proteins: POLRMT (P), TFAM (T), and TFB2M (B2), or TFB2M mutants ((GS)_3_-loop/Y209A/SGG-AAA), in reaction buffer containing 50 mM Tris-acetate (pH 7.5), 100 mM sodium glutamate, 10 mM magnesium acetate, 1 mM DTT, and 0.5 mM TCEP. The reaction mixture was added to increasing concentrations of ATP (ranging from 4 µM to 1400 µM). Fluorescence intensity was recorded at each addition step using a Fluoromax-4 spectrofluorometer at 25°C, with measurements taken in the 350-420 nm range (6 nm bandwidth) and an excitation wavelength at 315 nm (2 nm bandwidth). A control experiment was performed under the same conditions using unmodified LSP. The average fluorescence intensity between 360-380 nm of the modified LSP reaction sample was subtracted from the unmodified LSP reaction sample and the corresponding D-P-T-B2 complex (without ATP) to obtain F_obs_. The F_obs_ was plotted against [ATP] and the resulting hyperbolic increase was fit to Equation 2 to estimate the ATP dissociation constant (K_D_).

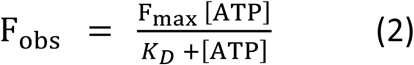

The *F*_max_ is the maximum increase in 2AP intensity upon ATP binding. Two independent experiments for each reaction sample have been performed.

### Fluorescence polarization assay

The K_D_ of the complex between Alexa488-Δ41 TFB2M and POLRMT-LSPDNA(-42to+9)-TFAM (P-D-T) complex was estimated using a fluorescence polarization assay on the Tecan Spark microplate instrument in 384-well black plates at 25°C. The 2:1:2 ratio of P-D-T was serially diluted and mixed with a fixed concentration of Alexa488-Δ41 TFB2M (30 nM) in reaction buffer (50 mM Tris-acetate, pH 7.5, 100 mM sodium glutamate, 10 mM magnesium acetate, 0.1 mg BSA, 0.5 mM TCEP) and incubated for 5 min at room temperature. Fluorescence polarization was measured after excitation at 488 nm (bandwidth 15 nm) and emission of 533 nm (bandwidth 20 nm). To estimate the K_D_ value of labeled TFB2M binding to P-D-T, fluorescence polarization values were plotted against the DNA concentration in the P-D-T complex and fitted to Equation 3, with the fitting errors reported. Experiments have been carried out in triplicates.

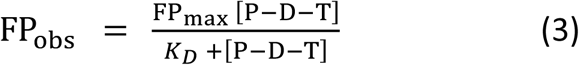

*FP*_obs_ and *FP*_max_ are the observed and final fluorescence polarization, respectively.

To estimate the binding affinity of the TFB2M mutants ((GS)_3_-loop, SGG-AAA and Y209A) to the P-D-T complex, we used a competition binding assay where the unlabeled TFB2M (mutants) were serially diluted in the reaction buffer and mixed with fixed concentrations of pre-mixed P-D-T (120:60:120 nM) and 30 nM Alexa488-TFB2M. The fluorescence polarization decreased as the unlabeled TFB2M increased in the mixture. To obtain the IC_50_ values, fluorescence polarization values were plotted against the concentration of unlabeled TFB2M, and the hyperbolic decrease was fitted to Equation 4. Two independent experiments for each titration reaction have been performed.

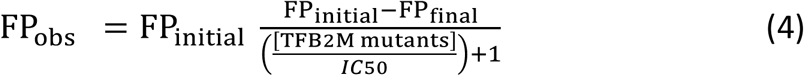

*FP*_obs_*, FP*_initial_ and *FP*_final_ are the observed, initial and final fluorescence polarization, respectively.

The Cheng–Prusoff – Equation 5 – gives the relationship between the inhibition constant (K_I_) and IC50 when a competitive inhibitor is present ^46^.

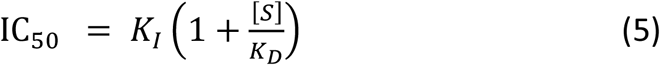

Where [*S*] is the concentration of Alexa488-TFB2M (30 nM), and K_D_ of TFB2M for P-D-T is 6.1 nM (Figure 6A) from the polarization experiment above.

### Labelling of TFB2M with Alexa488-C5-maleimide dye

Δ41 TFB2M (42-396 aa construct, 200 µl of 23 µM) was buffer-exchanged (50 mM HEPES pH 7.5, 350 mM NaCl and 0.5 mM TCEP) using a 2 ml 10 kDa cut-off AMICON ultra centrifugal concentrator (Merck Millipore Ltd.) and concentrated to ∼ 50 µM. Alexa fluoro488-C5-maleimide dye, 500 µM (Invitrogen) was added to 50 µM TFB2M (10:1 ratio) and incubated for ∼15 hours at 4°C in the dark. The unreacted fluorophore was removed using a 2 ml 10 kDa cut-off AMICON ultra centrifugal concentrator filter (Merck Millipore Ltd.) with 200 µl wash buffer (50 mM Tris acetate pH 7.5, 50 mM sodium glutamate) about 10 times until the flowthrough was colorless. The protein was analyzed by SDS PAGE to confirm the lack of free fluorophore. The efficiency of fluorophore incorporation in TFB2M was determined from the absorbance value of Alexa fluorophore and protein at 488 and 280 nm, respectively, and the molar extinction coefficient values of each using Equation 6:

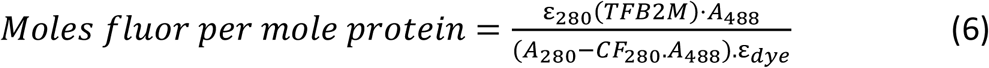

Where ε_280_(*TFB2M*) is 49,890 M^-1^.cm^-1^ and ε_dye_ for Alexa488 is 72,000 M^-1^.cm^-1^. The correction factor (*CF_280_*) is the ratio of the absorbances of Alexa fluoro488-C5-maleimide at 280 nm and 488 nm (0.11). The labelling efficiency of TFB2M was ∼76%. Finally, the functional activity of labeled TFB2M was verified using a transcription assay (Figure S6E).

### Assembly and characterization of h-mt IC3 and slipped IC3

For cryo-EM structural studies, Δ59 TFB2M was used as it purifies at a higher yield and forms a stable complex with POLRMT on the LSP promoter DNA, displaying a similar transcription activity as Δ20 TFB2M lacking only the mitochondrial localization signal (Figure S6F). The three component proteins of the human mitochondrial IC – POLRMT, TFAM and TFB2M – were individually expressed and purified as mentioned above. The h-mtRNAP open complex was prepared by incubating POLRMT, Δ59 TFB2M, TFAM and the biochemically optimized 56-bp bubble LSP promoter in a molar ratio of 1:1.5:1:1.1. in buffer A (30 mM Tris-HCl pH 8, 200 mM NaCl, 3 mM DTT, 10 mM MgCl_2_, 3% glycerol). The complex was purified by size-exclusion chromatography (SEC) in buffer B (30 mM Tris-HCl pH 8, 200 mM NaCl, 3 mM DTT, 10 mM MgCl_2_) using a Superdex 200 Increase 10/300 GL column connected to an AKTA PURE 25 FPLC system (GE Healthcare Life Sciences) maintained at 6°C. The complex eluted as a double peak, and the higher molecular-weight fractions had the complex of interest (Figures S1D and S1E). Stand-alone cuvette mode dynamic light scattering (DLS) experiments using a DynaPro Nanostar (Wyatt Technologies) on the SEC-purified complex provided a hydrodynamic radius of 6.1 nm with a polydispersity of 11.4% and an estimated molecular weight of ∼235 kDa (theoretical 232.4 kDa) (Figure S1F). Subsequently the fraction of interest was concentrated to 1.7 mg/ml (7 µM), aliquoted and stored in −80°C prior to further use. The concentration of the complex was determined through NanoDrop One UV-VIS spectrophotometer (Thermo Fisher) measurements with buffer B as blank. The isolated ternary open complex on the 56-mer bubble LSP promoter was incubated with GTP and 3’-dATP and supplemented with additional TFAM at a molar ratio of 1:35:35:1.1, respectively, to generate IC3; and incubated with pppGpG 2-mer RNA (TriLink BioTechnologies) and ATP at a molar ratio of 1:3:25 to prepare slipped IC3. For the DLS experiments, 10 µL of the sample was loaded into disposable cuvettes (Wyatt Technologies) and placed in the sample chamber maintained at 4°C. Each experiment consisted of 30 acquisitions and the obtained data was analyzed with Dynamics software (Version 7.10.0.23; Wyatt Technologies).

### Cryo-EM grid preparation and data collection

Cryo-EM grids of h-mtRNAP IC3 and slipped IC3 were prepared on Quantifoil R 1.2/1.3 holey carbon using a Vitrobot Mark IV-grid plunger (Thermo Fisher Scientific) and Leica EM GP (Leice Microsystems), respectively. The grids were glow-discharged for 45 s at 25 mA, with the chamber pressure set to 0.3 mbar (PELCO easiGlow Glow Discharge Cleaning System; Ted Pella). The grids were mounted in the sample chamber of Vitrobot Mark IV set at 4°C and 100% relative humidity and 8°C temperature and 95% relative humidity in Leica EM GP. Optimized grids for IC3 and slipped IC3 were obtained by spotting 3 µL of the sample each at 1.2 mg/ml (5 µM) and 0.3 mg/ml (1.4 µM) on Au300 grids. IC3 sample (in 30 mM Tris-HCl pH 8, 200 mM NaCl, 3 mM DTT, 10 mM MgCl_2_) was incubated on the grid for 5 seconds prior to double-sided blotting for 3.5 seconds at force 1. Slipped IC3 sample (in 30 mM Tris-HCl pH 8, 200 mM NaCl, 3 mM DTT, 10 mM MgCl_2_, 0.01% n-octyl-β-d-glucoside (BOG)) was incubated on the grid for 10 seconds prior to back-blotting for 12 seconds. After Plunge-freezing the grids in liquid ethane, grids were clipped and mounted on an in-house 200-keV Glacios cryo-transmission electron microscope with an autoloader and Falcon 3 direct electron detector for slipped IC3, and on the Glacios with post column Selectris energy filter and Falcon 4i direct electron detector for IC3 (Thermo Fisher Scientific). High resolution datasets for h-mtRNAP IC3 and slipped IC3 were collected on the Glacios using EPU software (versions 3.5.1.6034 & 2.9.0, respectively – Thermo Fisher Scientific). Electron microscopy data were recorded as movies in counting mode at a nominal magnification of x130.000 for IC3 and x150.000 for slipped IC3, yielding a pixel size of 0.9 Å and 0.97 Å, respectively. Both datasets were subjected to a total dose of 40 e/Å^2^ and were fractionated into 40 frames. Data collection parameters for both structures are listed in Table S1.

### Cryo-EM data processing

The workflows for IC3 and slipped IC3 structure determination are shown in Figures S5 and S7, respectively. Individual movie frames of both datasets were motion-corrected and aligned using MotionCor2^47^ as implemented in the Relion 3.1 and Relion 4 package for slipped IC3 and IC3, respectively. Motion-corrected data for IC3 was imported and processed in CryoSPARC and the contrast transfer function (CTF) parameters were estimated by patch CTF^48–50^. Contaminated micrographs, and micrographs with poor CTF fit resolution were removed from the processing path prior to picking initial particles with the blob picker tool in CryoSPARC with dimensions set to 100-160 Å. After extraction with a box size of 320^3^ pixels, the picked particles were cleaned by cycles of 2D classification. Once a clean particle set was obtained an ab-initio model was reconstituted and used as a template for re-picking and extracting particles using the same settings as above. The extracted particles were subjected to cycles of 2D classification (n=100) to clean up the dataset and subsequently an ab-initio reconstruction and hetero-refinement job (n=8) was run to separate the different populations from junk classes. Each obtained class was re-2D-classified and remaining 2D junk classes were excluded. The final cleaned particles for the five distinct populations were re-extracted and used in ab initio, heterogeneous refinement and non-uniform refinement jobs. The respective particle sets were used in global CTF refinement jobs prior to using them for non-uniform refinement and local refinement to obtain final models. Final models for State-I (IC3), State-II (IC3 – no TFAM), State-III (homodimer), State-IV (heterodimer) and State-V (free POLRMT) contained 175,482, 136,426, 46,546, 53,426 and 155,700 particles, respectively. Local resolution maps were calculated using the local resolution estimation tool in CryoSPARC and displayed in Chimera^51^ (Figure S5), whereas nominal resolutions for all maps were obtained using the FSC package in CryoSPARC.

The slipped IC3 dataset was processed using Relion3.1 (Figure S7) and the CTF parameters were estimated by CTFFIND-4^48^. Particles were automatically picked using reference-free Laplacian-of-Gaussian routine in Relion 3.1 with particle diameter set to 110-180 Å. Picked particles were subjected to cycles of 2D and 3D classifications to remove junk classes and obtain a homogeneous particle set. The best 3D class was used to calculate a gold-standard 3D auto-refined map and corresponding mask, and respective particles were repicked at the 3D auto-refined positions and re-extracted with a box size of 192^3^ pixels prior to further classification within its mask. Final 3D classification generated a distinct class, and respective particles were used to calculate the gold-standard auto-refined map, which was further improved by Bayesian polishing and CTF refinement. Both local resolution map – calculated using ResMap –, and the angular distribution were generated by Relion 3.1. The 3DFSC plots were calculated on the 3DFSC server (https://3dfsc.salk.edu/)^52^.

### Model building

Data processing yielded 3.4 Å and 3.15 Å density maps for IC3 (State-I – POLRMT/TFAM/TFB2M/DNA/RNA/GTP) and slipped IC3, respectively. In addition, the multiple states observed in the IC3 dataset yielded cryo-EM density maps calculated at 3.5 Å, 4.2 Å, 4.2 Å and 3.6 Å for state-II - POLRMT/TFB2M/DNA/RNA/GTP (IC3 – no TFAM); State-III - a homodimer of State-II interacting via the POLRMT PPR domain; State-IV a heterodimer of State-II with a POLRMT dimerized at their PPR domains; and State-V - POLRMT monomers, respectively. The obtained cryo-EM density maps were used to fit the atomic models for the respective structures. The previously published structure of the h-mtRNAP IC (PDB: 6ERP) was used as an initial model to build the models of the h-mtRNAP IC3, slipped structure and state-II (IC3 – no TFAM) after manually docking into the respective map using USCF Chimera. In addition, the Alphafold prediction of POLRMT (UniProt ID: O00411) and TFB2M (UniProt ID: Q9H5Q4) were used for modelling the two flexible helices between the tether-helix and PPR domain (residues 159 to 199) in POLRMT and the previously unidentified TFB2M loop (residues 268 to 294) in TFB2M. Model building for State-I (IC3), slipped IC3, and State-II (IC3 – no TFAM) were carried out manually using COOT and were coupled with iterative rounds of real-space structure refinement using Phenix 1.19.2^53,54^. No models were reconstructed for State-III and State-IV as the cryo-EM maps were of insufficient quality. All structure figures were generated using PyMol (https://pymol.org/2/), Chimera and ChimeraX^51,55^. Videos were generated using Chimera.

---

**Figure S1.**
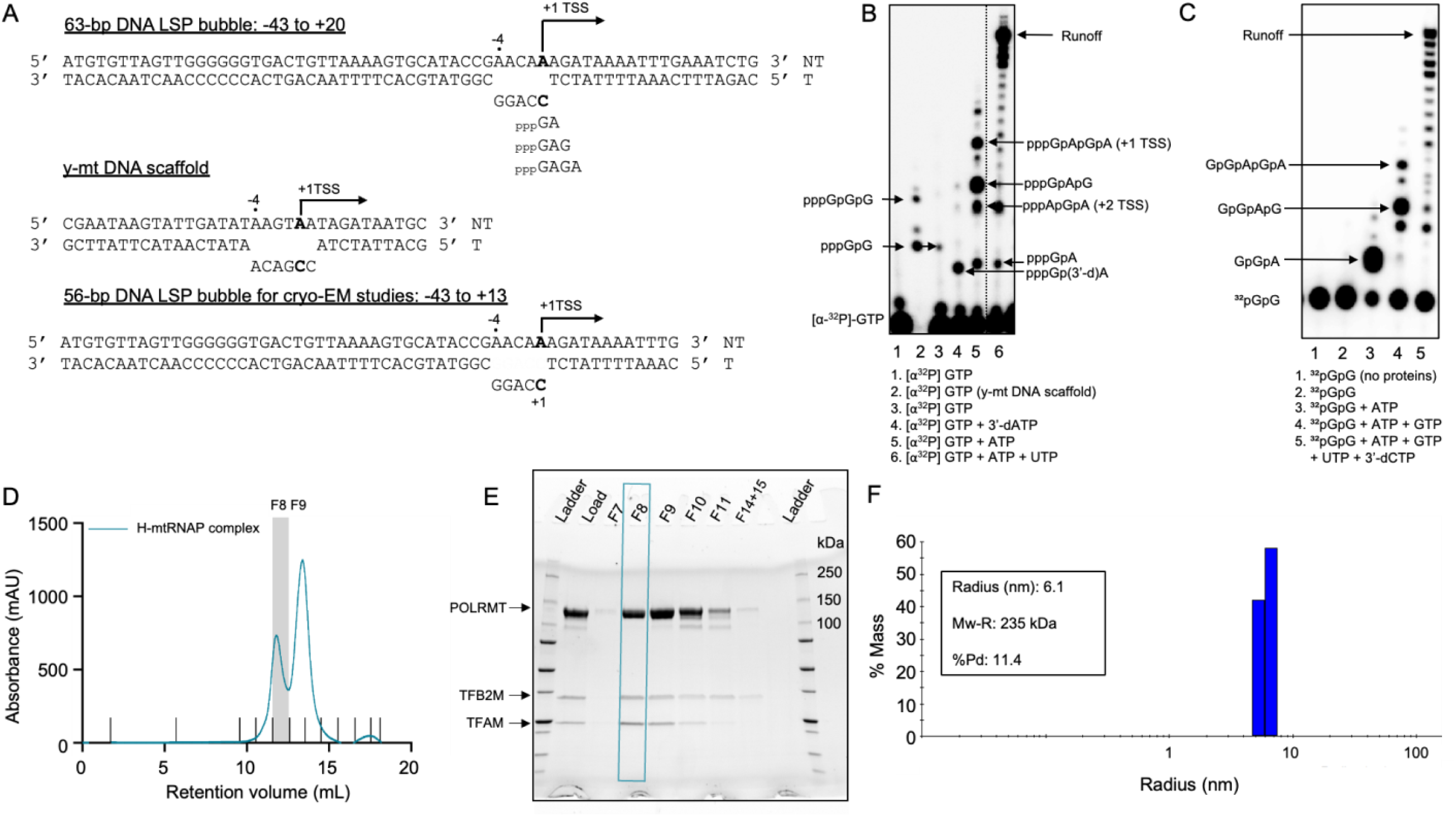
DNA scaffolds and biochemical/biophysical characterization of h-mtRNAP IC. (A) LSP bubbles used for *in vitro* transcription assays and cryo-EM studies. (B) Transcription reactions on 63-bp DNA LSP bubble with POLRMT, TFAM, and TFB2M in the presence of different NTP combinations. (C) Transcription reactions (same experimental conditions as in panel B) with [a^32^P]-GpG instead of [a^32^P]GTP. (D) Size-exclusion chromatography (SEC) profile of the complex between POLRMT, TFAM, and Δ59 TFB2M on the 56-bp LSP bubble (-4 to +1). The area in gray corresponds to fraction 8 (F8) which was used in cryo-EM structural studies. (E) SDS-PAGE of the eluted fractions from SEC in panel (D). The marked area corresponds to F8. (F) DLS profile of F8 indicated a monodisperse complex with hydrodynamics radius and estimated M.W. close to the theoretical value of the complex (232.4 kDa).

**Figure S2.**
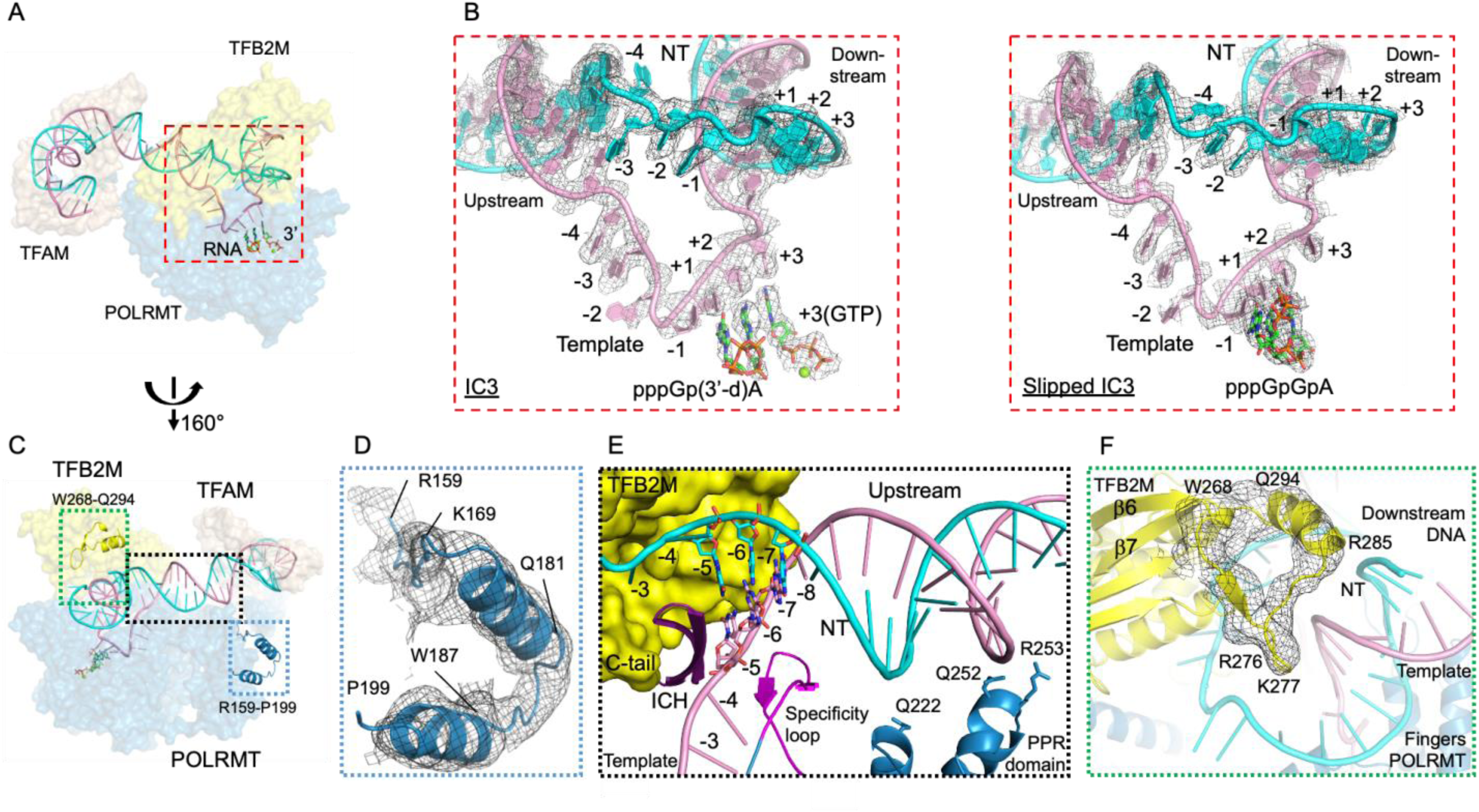
Structures of IC3 and slipped IC3 reveal fully resolved transcription bubble and previously unidentified regions. (A) Overview figure of IC3 with DNA tracks surrounded by protein shown in transparent surface representation (TFAM – brown; TFB2M – yellow; POLRMT – blue). (B) Zoomed view of the area outlined by the dashed red box in panel (A). B-sharpened experimental density maps define the fully resolved transcription bubbles in IC3 (left) and slipped IC3 (right) (non-template – cyan; template – pink; RNA – green by heteroatom). The contour levels of the density maps are 6s and 2s for IC3 and slipped IC3, respectively. (C) Overview figure of IC3 rotated 160° around y-axis from view in panel (A). The boxed regions are zoomed in subsequent panels. (D) Zoomed view of the previously unidentified region between the tether helix and PPR domain of POLRMT (R159-P199) modelled in the unsharpened density map shown at 7σ. (E) Zoomed view of the upstream region of the DNA, stabilized by the PPR domain, intercalating hairpin (ICH – purple), specificity loop (magenta) of POLRMT and TFB2M. (F) Zoomed view of the previously unidentified TFB2M loop (yellow) between β6 and β7 modelled in the unsharpened density map shown at 7σ.

**Figure S3.**
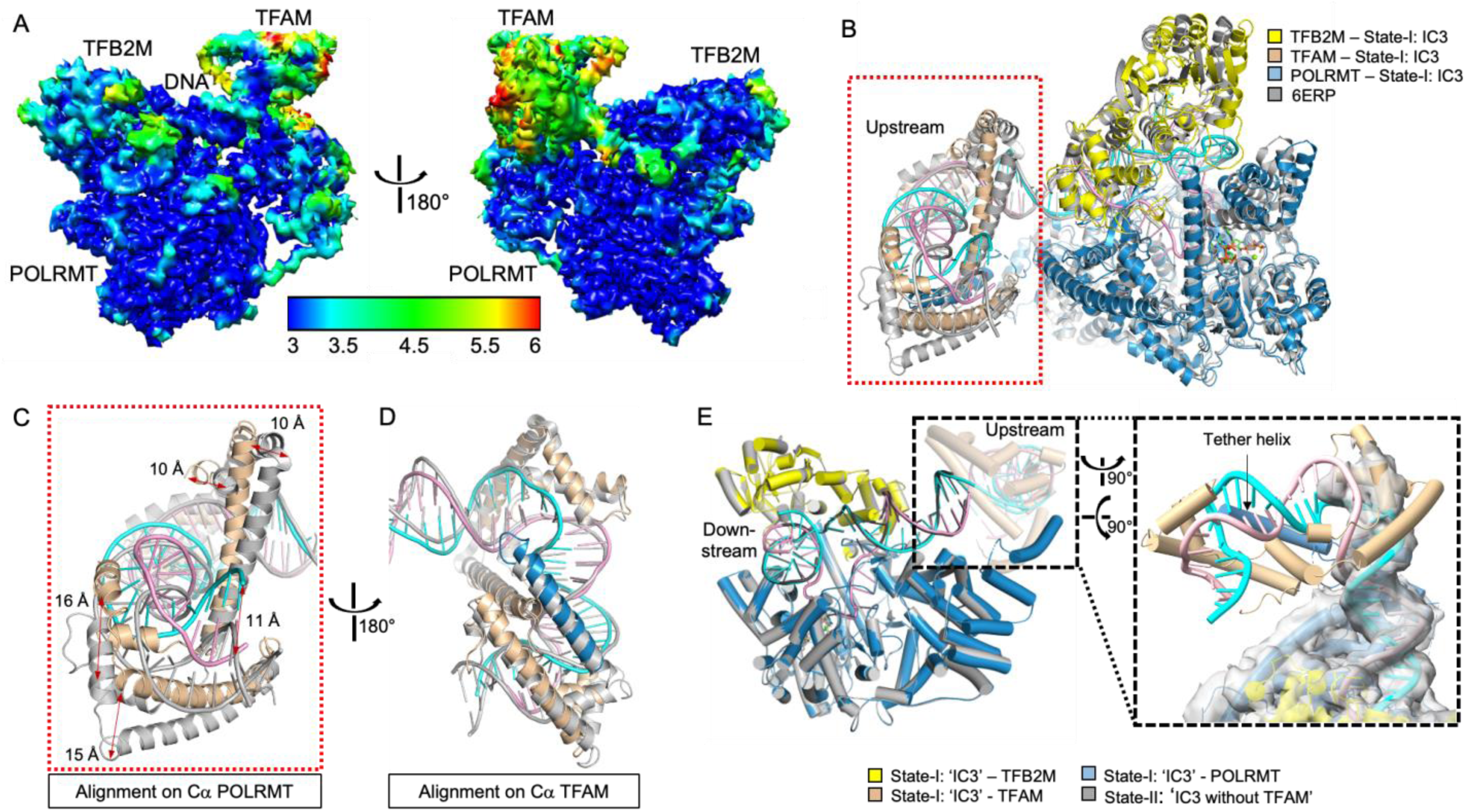
Comparison of h-mtRNAP transcription initiation complex structures. (A) Local resolution density map of h-mtRNAP IC3 generated by CryoSPARC (FSC threshold: 0.143) and surface colored in Chimera. (B) POLRMT Cα superposition of IC3 (color) with the crystal structure (gray – PDB: 6ERP). A total of 863 POLRMT Cα atoms were superimposed with an RMSD of ∼1 Å. (C) Zoomed view of the area outlined by the dashed red box in panel (B) highlighting the ∼15 Å shift in the position of TFAM between the cryo-EM and crystal structure (aligned on POLRMT Cαs). (D) View of panel (C) rotated 180° around y-axis. TFAM Ca superposition of IC3 (color) with the crystal structure (gray – PDB: 6ERP) shows negligible differences. A total of 183 TFAM Ca atoms were superimposed with an RMSD of ∼3 Å. (E) Left: POLRMT Cα superposition of ‘IC3’ (State-I – color) and ‘IC3 without TFAM’ (State-II – gray) show a highly superimposable core structure (849 Cα atoms with an RMSD of 0.3 Å) regardless of the presence of TFAM. TFAM and the extra resolved DNA in ‘IC3’ (State-I) is shown in transparent. Right: Zoomed view of the area outlined by the dashed black box rotated at 90° around the x- and z-axis. The ‘IC3’ structure (State-I) fitted in the unsharpened experimental density map of ‘IC3 without TFAM’ (State-II) show a straight trajectory of the upstream DNA in the absence of TFAM rather than bent track observed with TFAM. The unsharpened map is shown at 5σ.

**Figure S4.**
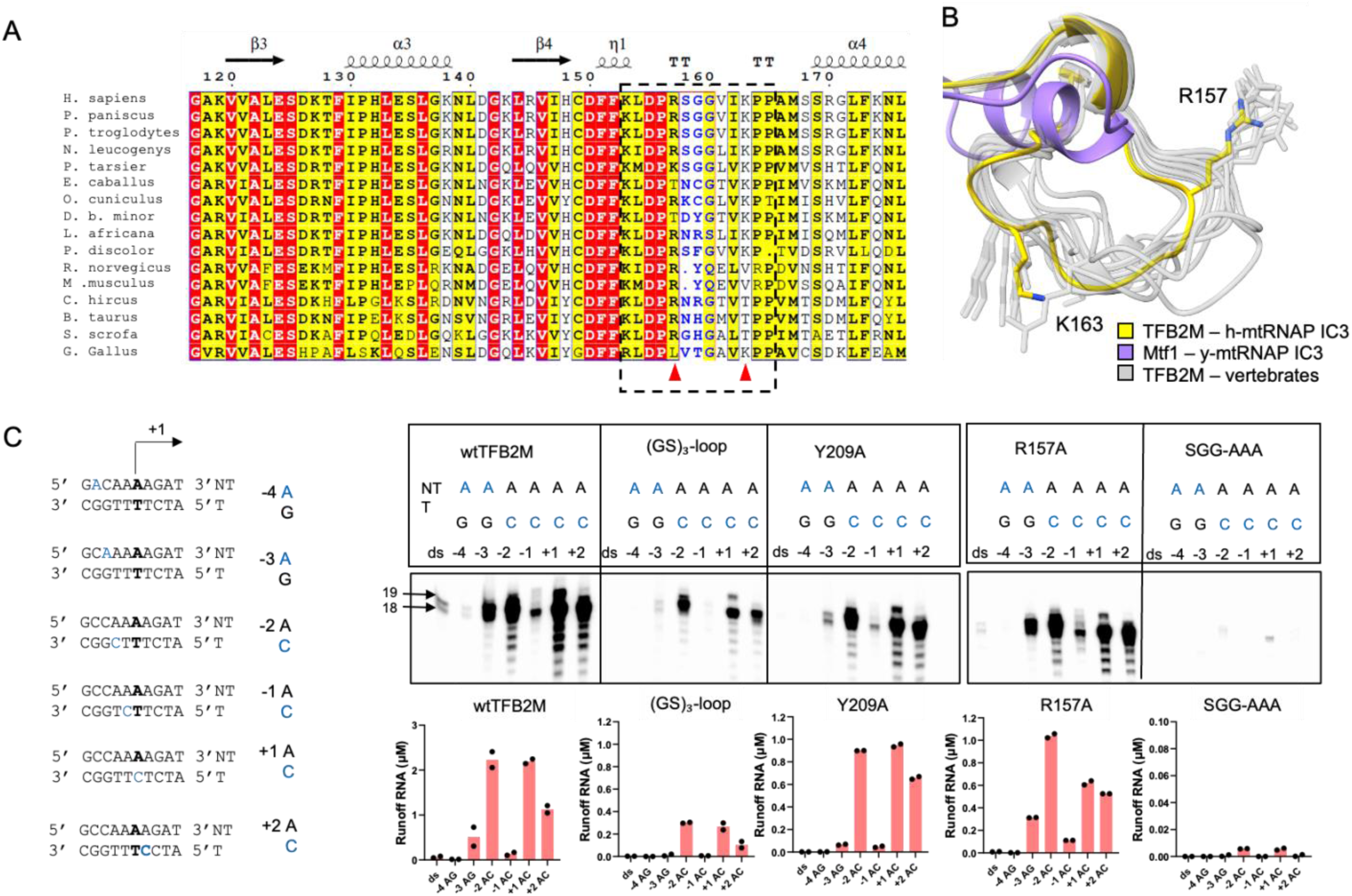
The structurally conserved NT-stabilizing loop in vertebrate TFB2Ms and effects of POLRMT, TFB2M and promoter mutations on transcription efficiency. (A) Sequence alignment of TFB2M across different vertebrates. The black dotted box indicates the start (K153) and end (P165) of the NT-stabilizing loop. The red triangles indicate R157 and K163 shown in panel (B). (B) Superposition of Mtf1 and the vertebrate TFB2Ms from panel (A) (generated by AlphaFold) suggesting that the NT-stabilizing loop is structurally conserved among vertebrates yet is absent in the yeast homolog. Mtf1 of y-mtRNAP IC3 is shown in purple, TFB2M of h-mtRNAP IC3 in yellow and AlphaFold generated models are in light gray. (C) LSP DNA sequence (-43 to +20) depicting mismatches from −4 to +2 (left). The +1 TSS site is highlighted in bold, and each mutation position is represented in blue. Transcription runoff assays show the −2, +1 and +2 mismatched positions partly rescue the transcription activity of TFB2M mutants (R157A, Y209A, and the (GS)_3_-loop) but transcription activity of the SGG-AAA mutant cannot be rescued by any of the mismatches. Quantification of the runoff products is shown in the graphs (bottom).

**Figure S5.**
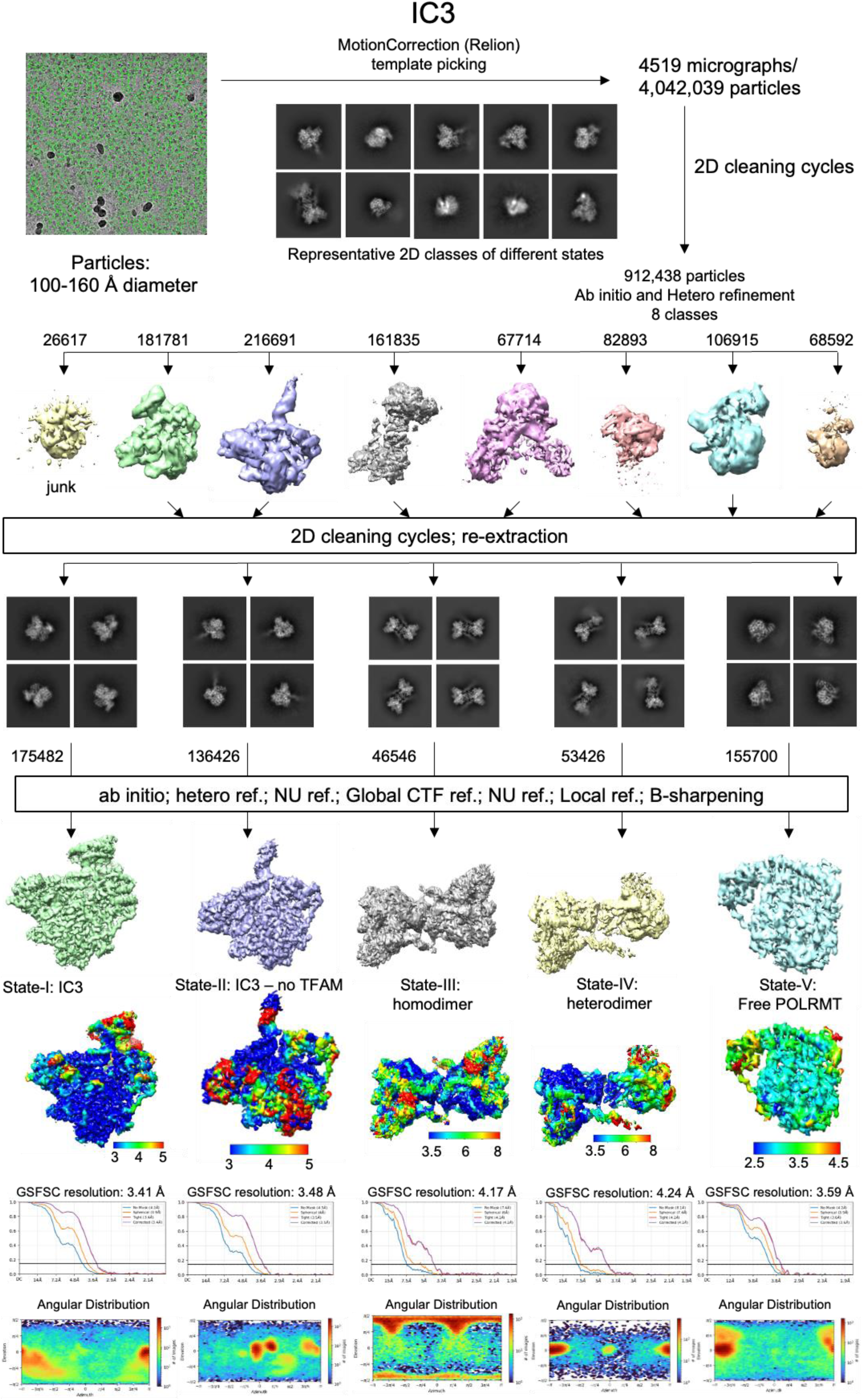
Cryo-EM image processing flowchart and density maps of IC3 (State-I), IC3 without TFAM (State-II), homodimer of State-II (State-III), heterodimer of State-II with POLRMT (State-IV) and free POLRMT (State-V). Processing flowchart of all states for obtaining the final maps. Five distinct populations were observed in the same dataset, labelled State-I – V. All jobs were processed in CryoSPARC unless differently mentioned. The local resolution maps, GSFSC curve of the final maps and angular distribution are shown.

**Figure S6.**
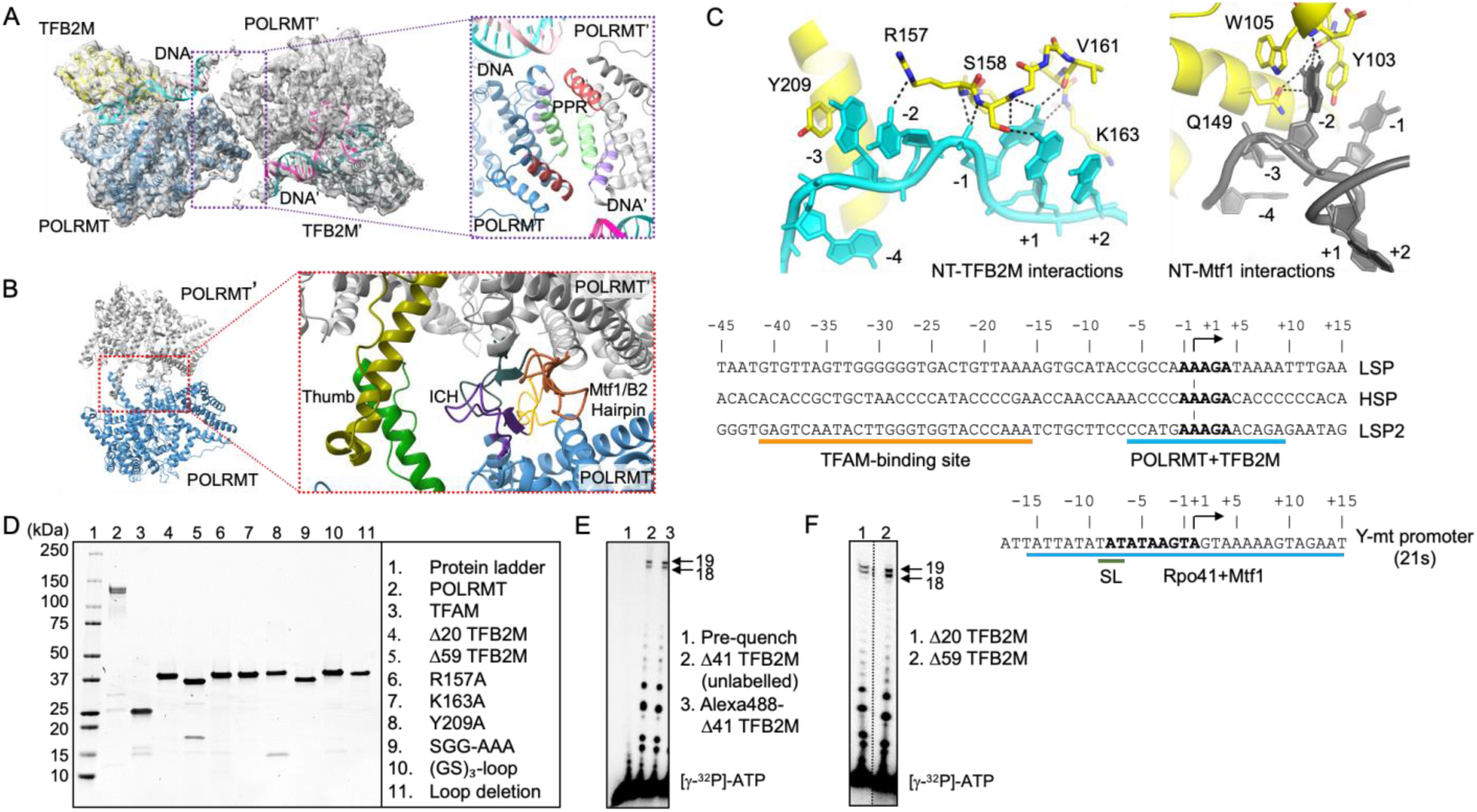
Single-subunit RNAP promoters and structural features stabilizing the non-template and dimer interfaces. (A) The PPR-domain-interfaced homodimer observed in the present study. Copy 1 (TFB2M – yellow; POLRMT – blue; non-template – cyan; template – pink) and copy 2 (TFB2M’ – dark gray; POLRMT’ – gray; non-template’ – slate; template’ – deep pink) are shown in the unsharpened experimental density map at 4σ. The purple dashed box highlights the dimerization at the PPR domain. The same helices from respective copies are colored alike. (B) The 7S RNA-mediated POLRMT homodimer (PDB: 7PZP). POLRMT copy 1 is shown in blue, POLRMT copy 2 (POLRMT’) in gray. The red dashed box highlights the dimer interface at the Mtf1/B2 hairpin (orange & brown), intercalating hairpin (ICH – purple & dark gray) and thumb (green & olive) subdomains. (C) Left: The NT-stabilizing loop of TFB2M and its base-specific interactions with −1 non-template in h-mtRNAP IC3. Right: Y103 and W105 of Mtf1 recognize the −2 non-template in y-mtRNAP IC3 (PDB: 6YMW). Bottom: Promoter sequences of human (LSP, HSP, LSP2) and yeast (21s) mitochondrial DNA (non-template strand). Conserved nucleotides are highlighted in bold. Yeast promoters are conserved from positions −8 to +1, human promoters are conserved from −1 to +4. Orange and blue indicate the binding sites for TFAM and POLRMT+TFB2M in humans, and Rpo41+Mtf1 in yeast, respectively. (D) SDS-PAGE gel shows the purified proteins used in the current study. (E) Transcription runoff assay confirms that the Alexa-488 labels on TFB2M do not impact the transcription activity. (F) Transcription runoff assay verifies that the TFB2M constructs have similar transcription activity.

**Figure S7.**
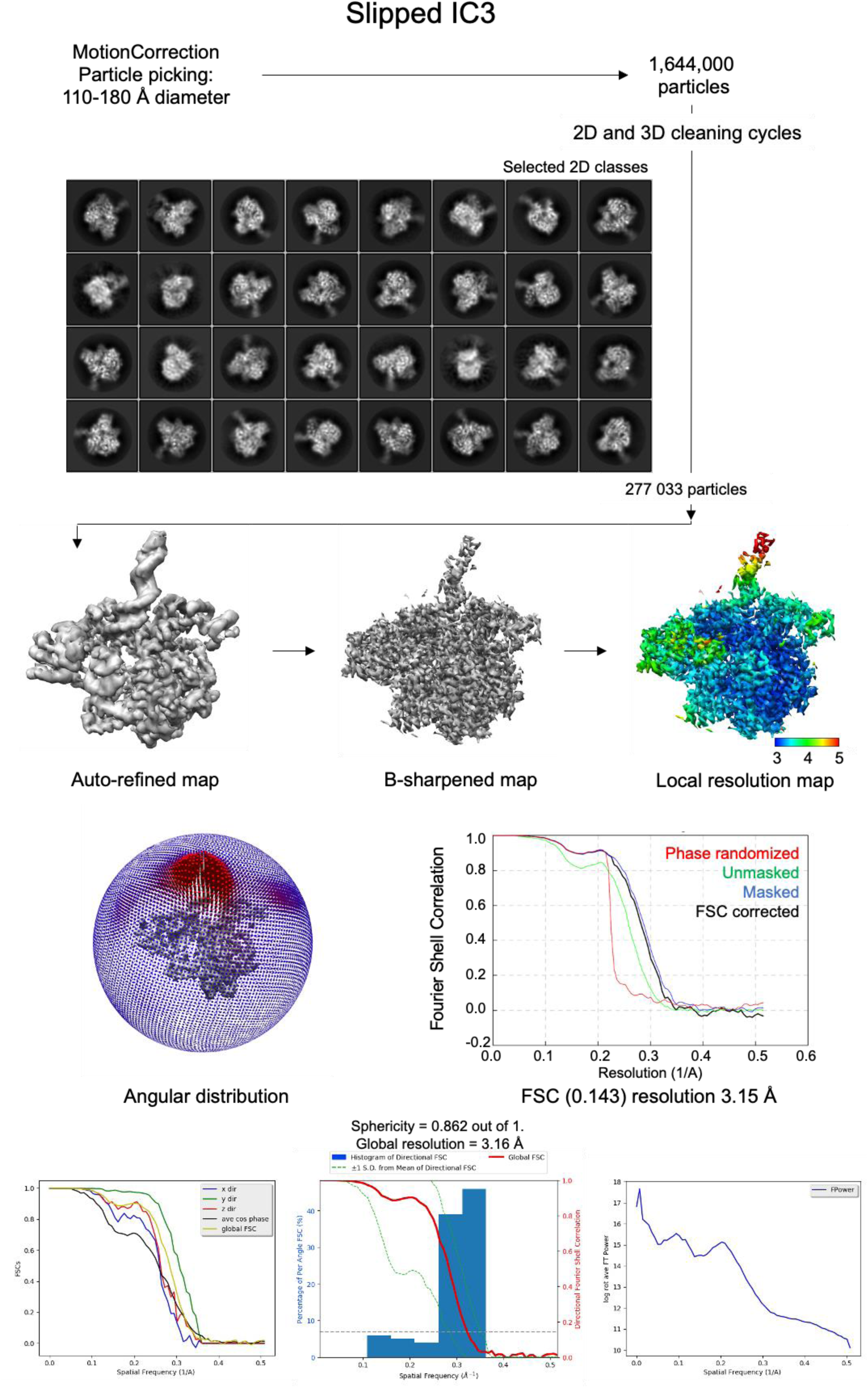
Cryo-EM image processing flowchart and density map quality of slipped IC3. Processing flowchart of h-mtRNAP slipped IC3 for obtaining the final map. All jobs were carried out using Relion. After B-sharpening, the final map was calculated at 3.15 Å. The local resolution map, angular distribution and FSC curve of the final map are shown. 3D FSC plots (bottom) for the cryo-EM map were calculated with https://3dfsc.salk.edu/.

**Video S1: The Human Mitochondrial Transcription Initiation Complex and the structural reorganization associated with to the opening and closing of the fingers domain**.

**00:00**: The animation shows an overview of the human mitochondrial transcription initiation complex.

**00:24**: In the closed conformation, the 3’ end of the nascent 3-mer RNA is positioned at the N-site of POLRMT. As RNA slippage occurs, the N-site becomes vacant, the fingers domain transitions to the open conformation, and residue Y999 moves into the N-site.

**00:34**: After NTP incorporation, Y999 shifts from the N-site, inducing a conformational switch that returns the fingers domain to the closed conformation.

**00:49**: In the closed state, the non-template (NT) strand is stabilized by the NT-stabilizing loop, Y209 of TFB2M, and W1026 of POLRMT. The opening of the fingers domain induces a structural rearrangement in the NT strand, causing a one-nucleotide shift in its interactions with TFB2M.

**00:59**: As the fingers domain transitions back to the closed conformation, the NT strand reverts to its previous state, reversing the one-nucleotide shift in its interactions with TFB2M. This cycle of fingers domain opening and closing, coupled with NT strand reorganization, repeats with each successive nucleotide addition.

Key Resources Table

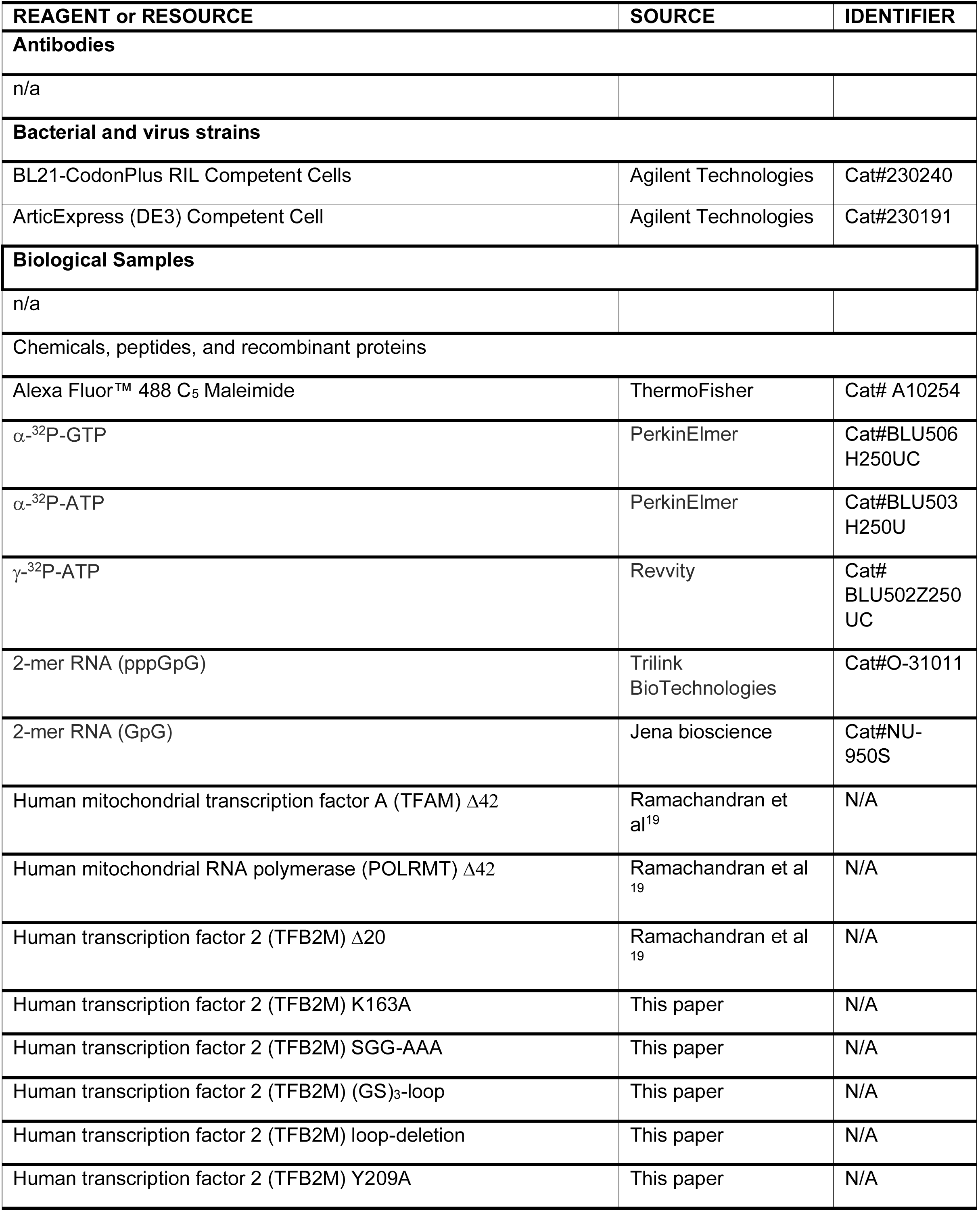
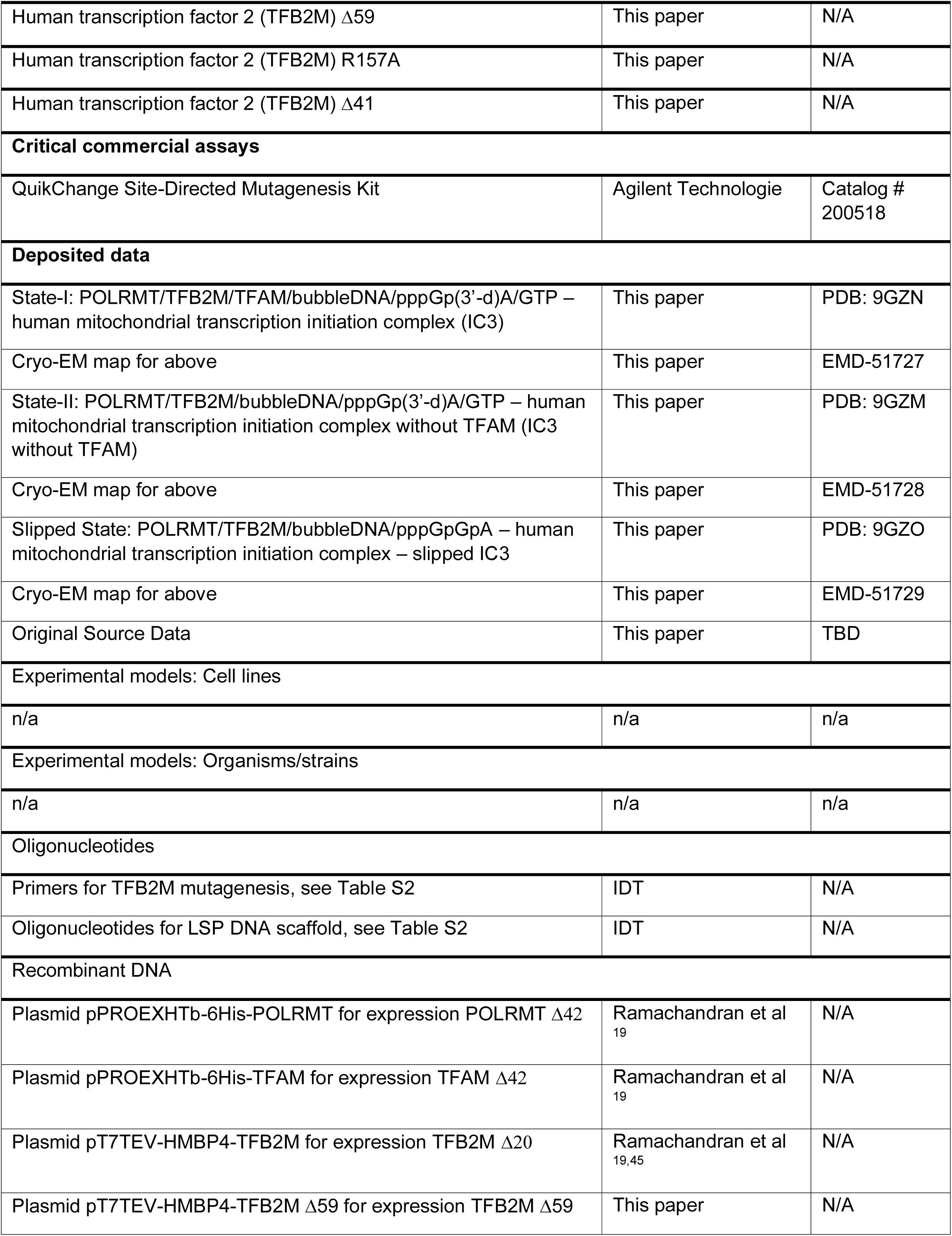
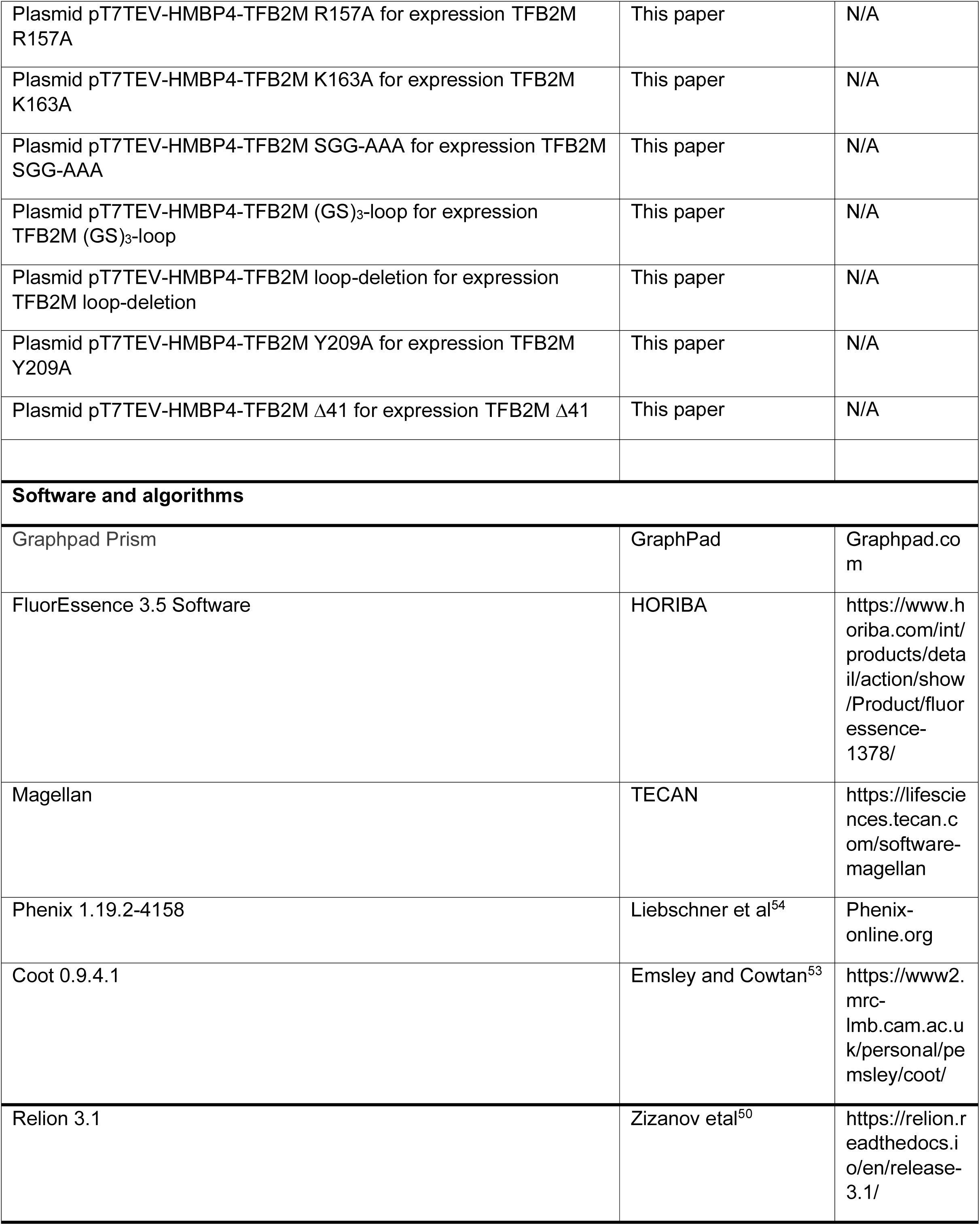
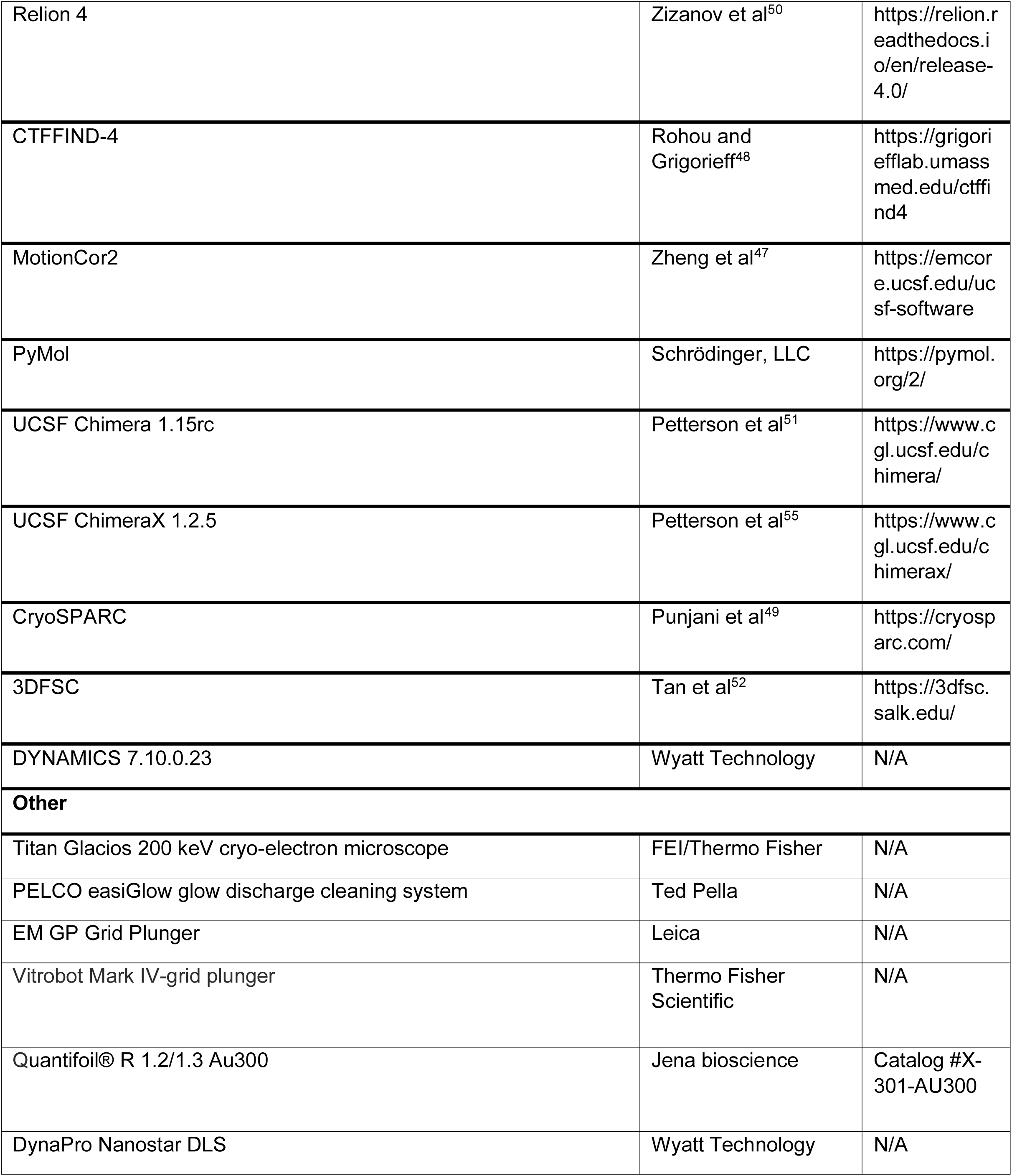

**Table S1.** Single Particle Cryo-EM data collection, refinement, and validation statistics.

|  | IC3<br>(POLRMT;TFB2M;TFAM<br>DNA;<br>pppGp(3'-d)A;GTP) | IC3 without TFAM<br>(POLRMT;TFB2M;<br>DNA;<br>pppGp(3'-d)A;GTP) | Slipped IC3<br>(POLRMT;TFB2M;<br>DNA;<br>pppGpGpA) |
| --- | --- | --- | --- |
|  | PDB 9GZM<br>EMD-51727 | PDB 9GZN<br>EMD-51728 | PDB 9GZO<br>EMD-51729 |
| **Data collection and processing** |  |  |  |
| Grid type | Quantifoil R1.2/1.3<br>Au300 | Quantifoil R1.2/1.3<br>Au300 | Quantifoil R1.2/1.3<br>Au300 |
| Microscope/detector | TFS Glacios/Falcon IV | TFS Glacios/Falcon IV | TFS Glacios/Falcon III |
| Magnification | 130,000x | 130,000x | 150,000x |
| Voltage (kV) | 200 | 200 | 200 |
| Electron exposure<br>(e<sup>-</sup>/Å<sup>2</sup>) | 40 | 40 | 40 |
| Defocus range (μm) | -0.6 to -1.8 | -0.6 to -1.8 | -0.6 to -1.8 |
| Pixel size (Å) | 0.9 | 0.9 | 0.97 |
| Symmetry imposed | C1 | C1 | C1 |
| Initial particle images<br>(no.) | 912,438 | 912,438 | 1,644,000 |
| Final particle images<br>(no.) | 175,482 | 136,426 | 277,033 |
| Map resolution (Å) | 3.41 | 3.48 | 3.15 |
| FSC threshold | 0.143 | 0.143 | 0.143 |
| Map resolution range<br>(Å) | 3.06-6.11 | 3.15-7.66 | 3-6.8 |
| **Refinement** |  |  |  |
| Initial model used<br>(PDB code) | 6ERP | 6ERP | 6ERP |
| Model resolution (Å) | 3.2 | 3.3 | 2.9 |
| FSC threshold | 0.143 | 0.143 | 0.143 |
| Model resolution<br>range (Å) | 100 to 3.2 | 100 to 3.3 | 100 to 2.9 |
| Map sharpening B<br>factor (Å) | -116 | -112 | -80 |
| **Model composition** |  |  |  |
| Non-hydrogen atoms | 14771 | 11633 | 11445 |
| Protein residues | 1583 | 1326 | 1285 |
| **B factors (Å)** |  |  |  |
| Protein | 75 | 67.74 | 71.78 |
| Nucleid acid and NTP | 145.64 | 102.78 | 128.38 |
| **R.m.s deviations** |  |  |  |
| Bond lengths (Å) | 0.008 | 0.006 | 0.007 |
| Bond angles (°) | 1.382 | 1.260 | 1.232 |
| **Validation** |  |  |  |
| Molprobity score | 1.59 | 1.58 | 1.61 |
| Clashscore | 7.78 | 6.09 | 8.08 |
| Poor rotamers (%) | 0.88 | 0.62 | 0.56 |
| **Ramachandran plot** |  |  |  |
| Favored (%) | 97.07 | 96.29 | 97.01 |
| Allowed (%) | 2.74 | 3.56 | 2.99 |
| Disallowed (%) | 0.19 | 0.15 | 0.00 |

**Table S2.**
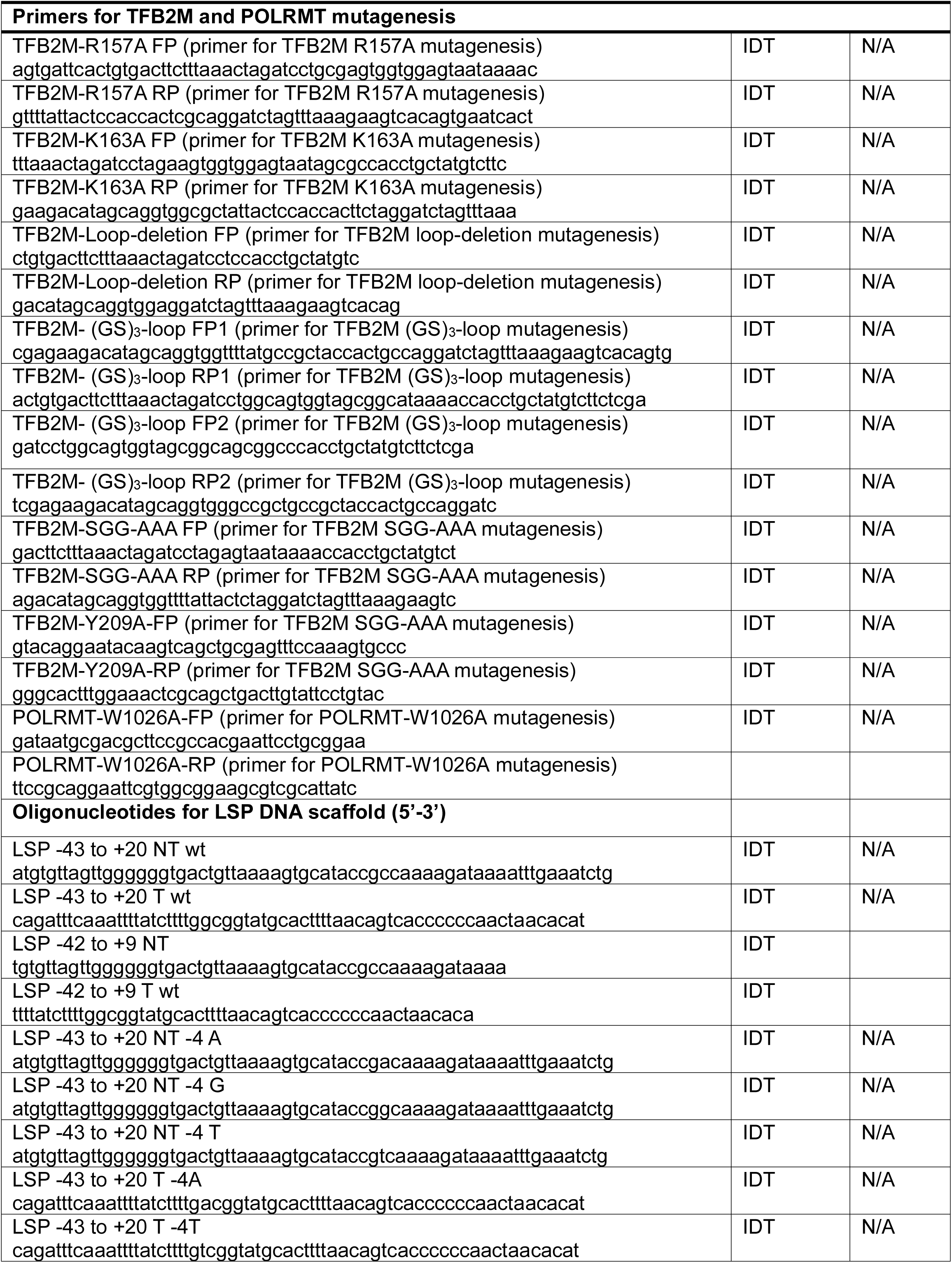
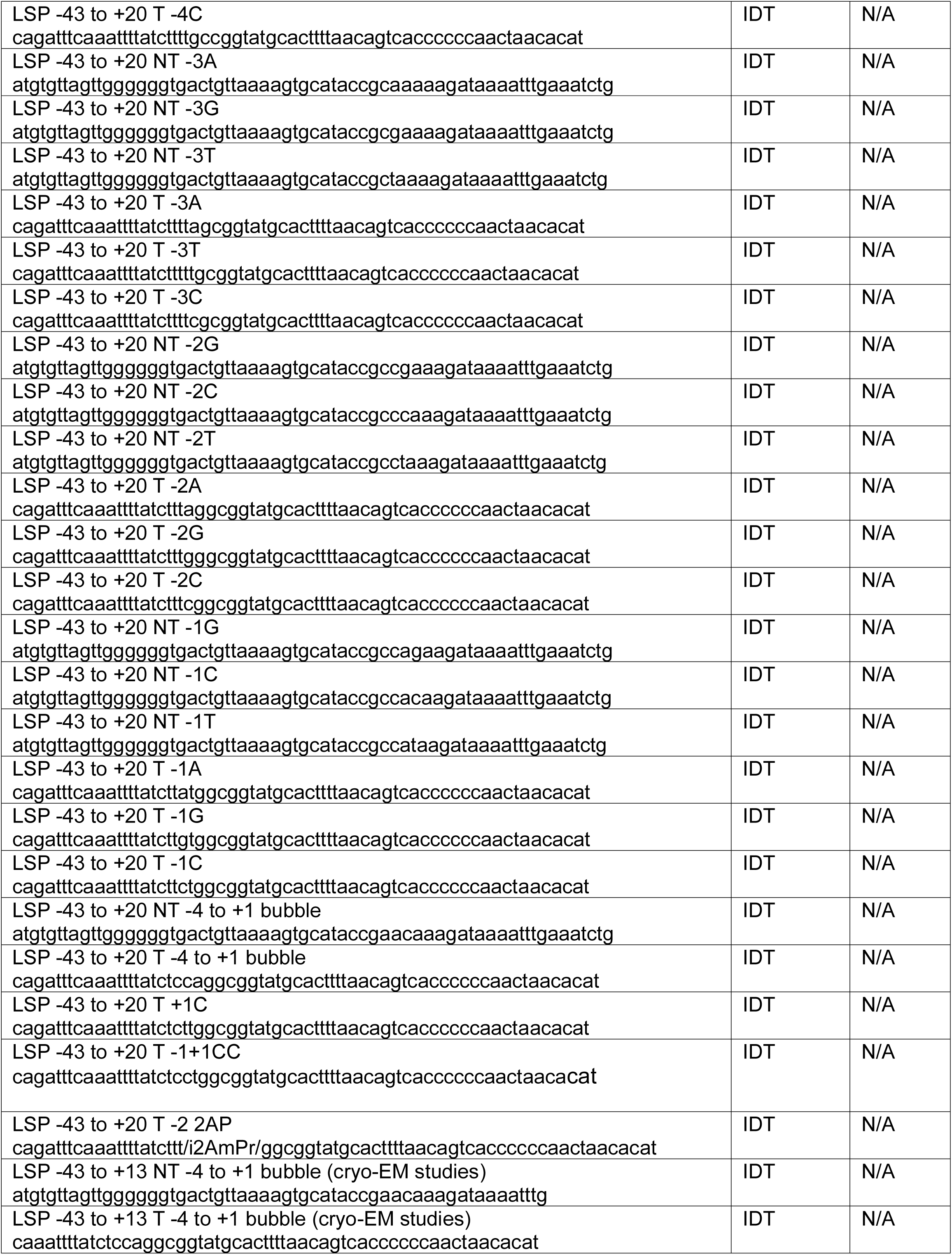
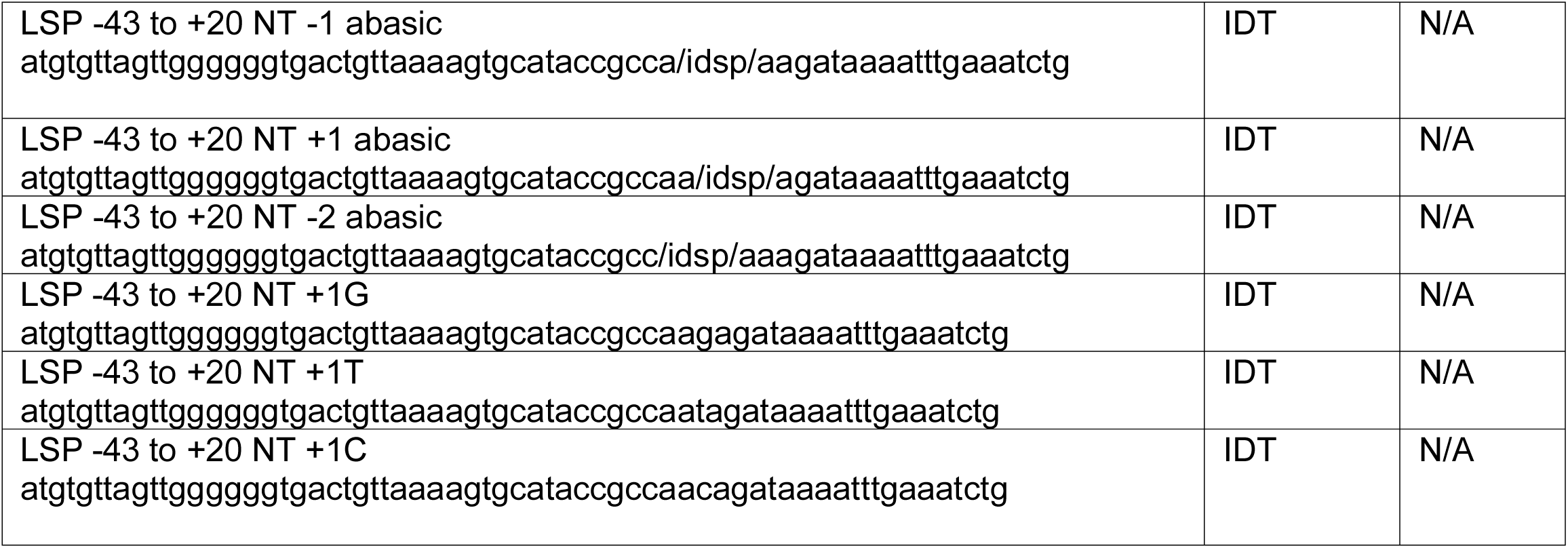
Oligonucleotides.

